# Pharmacological interrogation of crosstalk between TLR signaling and stress granules reveals a compound with antiviral effect

**DOI:** 10.1101/2025.04.11.648452

**Authors:** Aditi, Prem Prasad Lamichhane, Xuping Xie, Parimal Samir

## Abstract

Stress granules (SG) are cytoplasmic membraneless compartments that regulate cellular stress responses and have been implicated in antiviral defense against Influenza A Virus (IAV). SG stabilization has been reported to be a viable strategy to develop antiviral drugs. To expand the repertoire of SG-modulating compounds, we performed a targeted pharmacological screen focusing on inhibitors of TLR signaling pathway, based on our previous work demonstrating an antagonistic relationship between SGs and TLR signaling. Using the synthetic dsRNA analog polyinosinic:polycytidylic acid (poly(I:C)) to induce SGs in A549 human lung cancer cell line, we screened a panel of TLR signaling pathway inhibitors for their effect on SG. We developed a robust image analysis pipeline utilizing ilastik for pixel classification and CellProfiler for object quantification, enabling high-throughput analysis of SG alterations. This screen identified multiple small molecules that destabilize SGs, including inhibitors of RIPK3, TBK1, PERK, IRAK1/4, and JNK1/2/3 and ERK1/2 kinases. Several inhibitors of NF-κB signaling also disrupted SG integrity. Most strikingly, PPM18, an NF-κB inhibitor, emerged as a dual-action compound, triggering spontaneous SG assembly via PERK-mediated eIF2α phosphorylation, while simultaneously suppressing poly(I:C)-induced SGs. Intriguingly, PPM18 exhibited potent IAV inhibition, but crucially, this antiviral activity was decoupled from SG formation as it robustly inhibited replication in *G3BP1^-/-^* cells which did not assemble SGs upon PPM18 treatment. Our work not only identifies PPM18 as a unique SG modulator and an antiviral agent but also challenges the prevailing belief linking SG assembly directly to antiviral activity as a general phenomenon. Our findings demonstrate that SG formation and antiviral activity can be functionally uncoupled, providing new insights into host-virus interactions. In summary, our results demonstrate feasibility of using SG screening for finding novel antiviral compounds while raising important questions about the role of SGs themselves in modulating host-virus interactions.

## Introduction

Stress granules (SG) are dynamic cytoplasmic mRNA-ribonucleoprotein complexes that are assembled in response to variety of stressors^1–3^. SGs play an integral role in maintaining homeostasis by protecting stressed cells from regulated cell death (RCD)^4–7^. The canonical SG assembly follows a well characterized pathway initiated by activation of stress sensing kinases, that includes EIF2AK1 (heme regulated inhibitor (HRI)), EIF2AK2 (protein kinase R (PKR)), EIF2AK3 (PKR-like endoplasmic reticulum protein kinase (PERK)), and EIF2AK4 (general control nonderepressible 2 (GCN2))^7^. These kinases sense different types of stress signals and activate the integrated stress response (ISR) by phosphorylating a translation initiation factor eIF2α. Phosphorylation of eIF2α inhibits cap-dependent translation initiation, leading to accumulation of a stalled translation initiation complexes that, together with SG scaffold proteins, mRNAs and RNA binding proteins, condense to assemble SGs. However, translation initiation inhibition alone is not sufficient as evidenced by inability of puromycin to assemble SGs^8^. Likewise, phosphorylation of eIF2α alone is not sufficient to trigger SG assembly^9^, suggesting additional regulatory steps between translation initiation inhibition and biomolecular condensation that forms SGs.

Beyond their role in stress adaptation, there is a strong body of evidence that shows SGs modulate many cellular activities, including antiviral innate immune response^10–14^. SGs promote type I interferon signaling in response to viral infections^15–17^, and PKR activation can result in SG assembly^16,18–21^. To counteract this, viruses have evolved mechanisms to interfere with SGs^15,22–24^. For example, NS1 protein of Influenza A Virus (IAV-NS1) interferes with PKR to inhibit signal transduction downstream of its activation^25^. IAV-NS1 has also been reported to bind to a SG scaffold protein named DDX3X to directly interfere with SG assembly^15,17^. Interestingly, forced induction of SGs by orthogonal application of physicochemical stressors such as sodium arsenite greatly attenuates IAV replication and amplifies type I interferon signaling^17^. These findings have led to a model in which SGs function as localized signaling hubs that increase local concentration of viral pathogen associated molecular patterns (PAMPs) in a small volume of cytoplasm of infected cells. Host pattern recognition receptors (PRRs) are recruited to these areas of increased PAMPs where they sense infection and induce a robust antiviral host response^12–14,26^. Although, some studies have challenged aspect of this model^27–29^, the antiviral role of SGs itself is widely accepted, with only a few exceptions. Notably, a recent study reported that a compound that stabilized SGs inhibited replication of respiratory syncytial virus (RSV) in cells and animal models^30^. Consequently, pharmacological induction or stabilization of SGs has emerged as a promising strategy to develop host-directed antiviral drugs.

Our previous work uncovered an unexpected link between the toll-like receptor (TLR) signaling pathway and SG dynamics^9^. We demonstrated that TLR signaling inhibits SG assembly in bone marrow derived macrophages (BMDMs) and even promotes rapid disassembly of pre-formed SGs. In a follow-up study, we found that TLR signaling mediated SG inhibition was ineffective in cells with weak activation of the IKK complex^31^. While many kinases within the TLR signaling pathways are ubiquitously expressed and known to be activated by cellular stress such as mitogen activated protein (MAP) kinases and stress activated MAP kinases (for example, JNK and P38), their roles in SG regulation during innate immune activation remain largely unexplored.

Given the established antiviral function of SGs and the emerging potential of SG stabilization as a therapeutic approach, we hypothesized that inhibition of kinases downstream of TLR signaling could modulate SG assembly and disassembly. To test this, we designed a pilot screen targeted at TLR signaling kinases to identify small molecules that can modulate SGs formed in response to PKR activation. Our screen revealed several compounds that modulate SG dynamics. PPM18, an inhibitor of IKK complex, emerged as the strongest inducer of spontaneous SGs and robustly inhibited IAV replication. Our findings suggest an intricate crosstalk between innate immune signaling and SG regulation that has not been previously appreciated and offers new insights into how SG dynamics might be regulated during infection, inflammation, and disease.

## Results

### Design of the screen and development of a machine learning assisted data analyses pipeline

To identify compounds that could stabilize SGs formed downstream of PKR activation, we utilized cytosolic delivery of synthetic dsRNA analog polyinosinic:polycytidylic acid (poly(I:C)) through transfection in human A549 lung cancer cell line (**Figure 1A**). A549 cells were selected for these experiments due to their widespread use to study biology of Influenza A Virus. To maximize throughput, we performed these experiments in 96-well plates. Since we were interested in compounds that stabilize SG, we first transfected cells with poly(I:C) and incubated them for 6 hours to allow SG assembly. We then added inhibitors for an empirically selected list of kinases known to be activated downstream of TLR signaling and incubated for 1 hour (**Supplementary Figure 1**)^32–34^. Given that intracellular effective concentrations of many kinase inhibitors in A549 cells remain largely unknown, we tested at least 4 concentrations for every inhibitor (**Supplementary table 1**). These concentrations ranged from at least 10 times the reported IC50 concentration to a 64-fold lower concentration. Whenever available, published effective concentration of kinase inhibitors in cells were included. We used G3BP1 as a marker to visualize SGs using immunofluorescence confocal microscopy. Automated acquisition of images resulted in over 10,000 images necessitating development of an automated image analysis workflow. While several such tools are available from commercial vendors and academic labs, we developed a workflow integrating two open-source tools iLASTIK^35^ (for training machine learning models for SG detection and generating probability maps) and CellProfiler^36^ (for processing probability maps and images). These tools were chosen for their accessibility, active user community and ease of distribution, ensuring that our workflow can be readily shared with broader research community. We manually validated our analysis workflow on a subset of images (**Figure 1B**).

**Figure 1:**
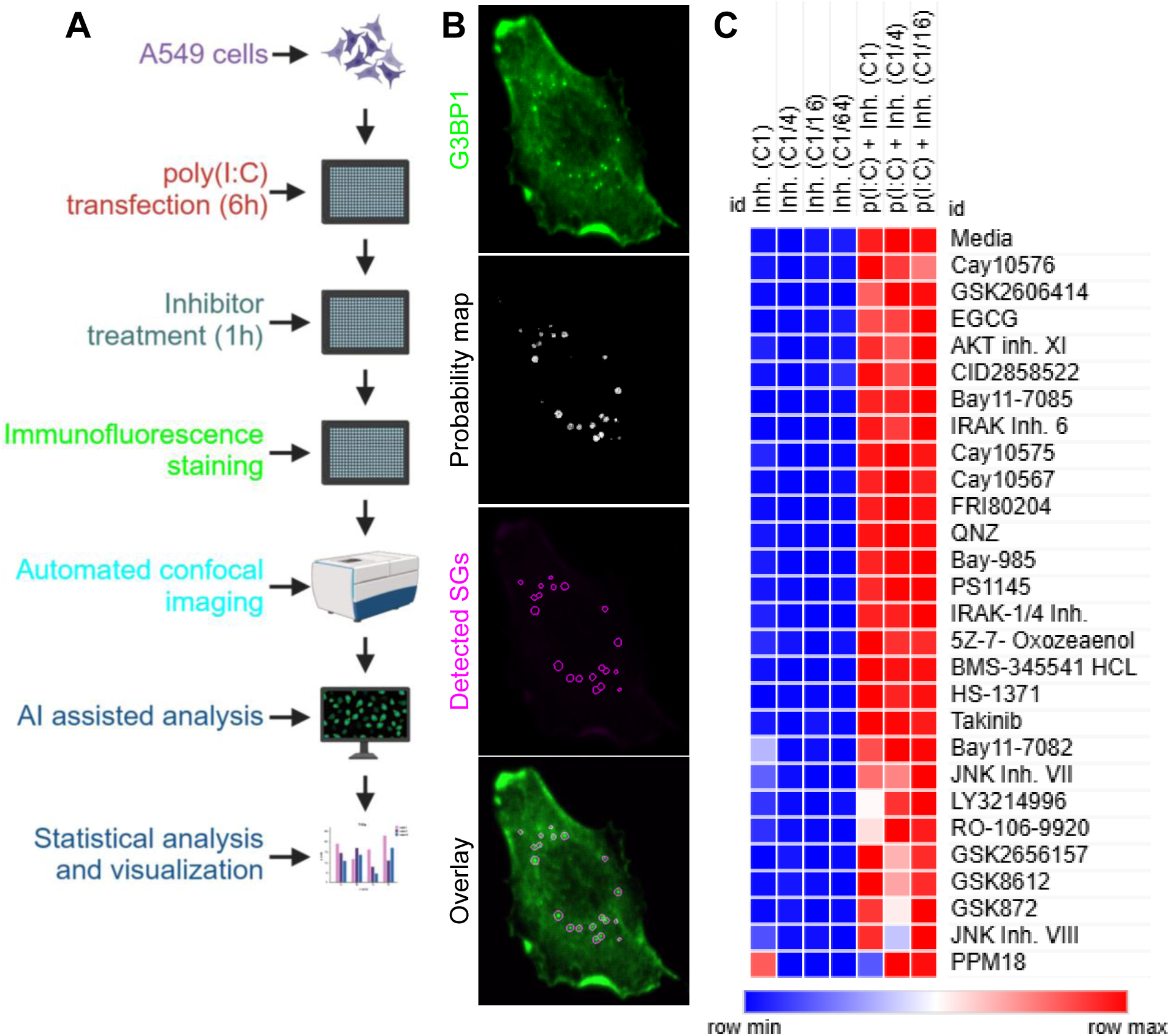
Screen design and data summary. **A**) Overview of screen design. A549 cells were transfected with polyinosinic:polycytidylic acid (p(I:C)) and incubated for 6 hours to allow assembly of SGs. Cells were then treated with inhibitors for 1 hours, fixed, immunostained and imaged. An artificial intelligence model was trained using iLASTIK and all images were batch processed to generate probability maps for SG detection. Probability maps and imaged were processed using CellProfiler to generate tables with SG quantification. **B**) An example of raw images and SGs identified using our workflow. Probability maps were generated using iLASTIK which was used as an input in a CellProfiler workflow to identify SGs. G3BP1 was used as the SG marker. DAPI was used to visualize nuclei. Identified SGs are marked in magenta. **C**) A heatmap of SG quantification results. Percent SG^+^ cells are reported. Color in heatmap is based on row minimum and maximum values (range = 0-100). Semi-supervised hierarchical clustering analysis on rows was used to organize results for tested compounds. Four concentrations of every inhibitor were used in a dilution series, with 4-fold dilution in each step. This resulted in a dynamic range of 64-fold concentration differences that were tested.

Our screen revealed several compounds that modulate SG dynamics, including unexpected instances where compounds induced SGs even in the absence of poly(I:C) transfection or destabilized SGs despite poly(I:C)-induced activation. These observations underscore the complex and context-dependent regulation of SG dynamics by innate immune signaling pathway kinases (**Figure 1C**). Our findings also suggest that SG regulation may be finetuned by different signaling events and highlights the need for further investigation into how innate immune responses and SG formation intersect to influence cellular outcomes in various pathological contexts.

### Inhibitors targeting kinases that are activated downstream of TLR activation alter SGs stability

Our screen revealed several unexpected connections between the TLR signaling pathway and SG regulation. TLRs are key sensors of the innate immune system that recognize exogenous PAMPs, and damage associated molecular patterns (DAMPs) to trigger a robust immune response. This signaling is primarily transduced through two adaptors, MYD88 and TRIF (TICAM1), which initiate downstream kinase cascades that modulate various cellular processes, including inflammatory gene expression and metabolic reprogramming. To explore how TLR signaling affects SG stability, we focused on empirically selected key kinases in the TLR pathway.

MYD88 assembles a multi-protein complex called the myddosome^37–41^ that relies on two protein kinases IRAK1 and IRAK4 (henceforth collectively called IRAK1/4) as catalytic engines for downstream signal transduction. Strikingly, treatment with either of the two small molecule inhibitors of IRAK1/4 (IRAK inhibitor 6 and IRAK-1/4 inhibitor) destabilized SGs formed downstream of dsRNA sensing (**Figure 2A-B**). This data suggests a novel role for IRAK1/4 in SG biology wherein it stabilizes SGs formed downstream of dsRNA induced PKR activation.

**Figure 2:**
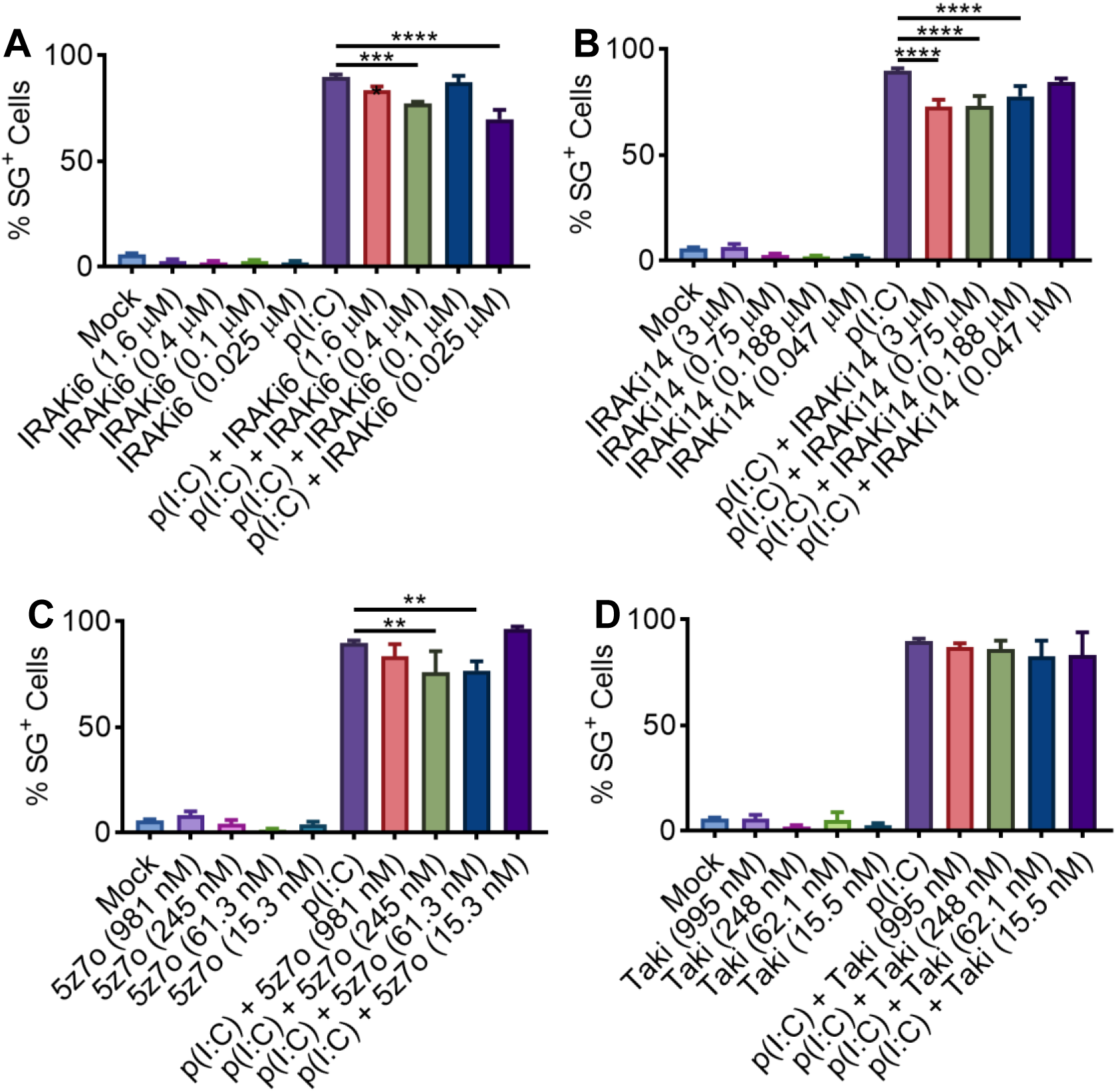
IRAK1/4 inhibitors destabilize SGs while TAK1 inhibitors have a differential effect. **A**) Quantification of SG^+^ cells after IRAK Inhibitor 6 treatment (IRAKi6). IRAK Inhibitor 6 inhibits kinase activity of IRAK4. **B**) Quantification of SG^+^ cells after IRAK-1/4 Inhibitor (IRAKi14) treatment. IRAK-1/4 Inhibitor can inhibit both IRAK1 and IRAK4 kinases. **C**) Quantification of SG^+^ cells after 5Z-7-Oxozeaenol (5z7o) treatment. 5Z-7-Oxozeaenol inhibits kinase activity of TAK1. **D)** Quantification of SG^+^ cells after Takinib (Taki) treatment. Takinib inhibits kinase activity of TAK1. One-way ANOVA was used to determine the statistical significance. \*\**p*-value<=0.01, \*\*\**p*-value<=0.001, and \*\*\*\**p*-value<=0.0001.

TRIF is the adapter protein activated by two specific TLRs (TLR3 and TLR4)^37–41^. TRIF mediated signaling activates TAK1 (MAP3K7) which integrates the TRIF and MYD88 arms of TLR signaling network. We used 5Z-7-Oxozeaenol and Takinib inhibitors to study the role of TAK1 in SG biology. Treatment with 5Z-7-Oxozeaenol destabilized SGs while Takinib treatment did not have an effect (**Figure 2C-D**). TBK1 is another kinase in the TLR signaling network that is activated through the TRIF arm. Therefore, we examined the role of TBK1 using two compounds named Bay-985 and GSK8612. Bay-985 did not affect SG stability, while treatment with GSK8612 destabilized SGs (**Figure 3A-B**). Additionally, treatment with a multi-kinase inhibitor named Cay10576 that inhibits TBK1, IKKβ and PLK1 also destabilized SGs (**Figure 3C**). Interestingly, Cay10575, which inhibits IKKβ and PLK1 kinases, did not affect SG stability suggesting that SG destabilizing activity of Cay10576 is likely due to its inhibition of TBK1 (**Figure 3D**). However, given that Bay-985 did not affect SGs, the connection between TBK1 and SGs needs further investigation using genetic knockout cells before drawing firm conclusions.

**Figure 3:**
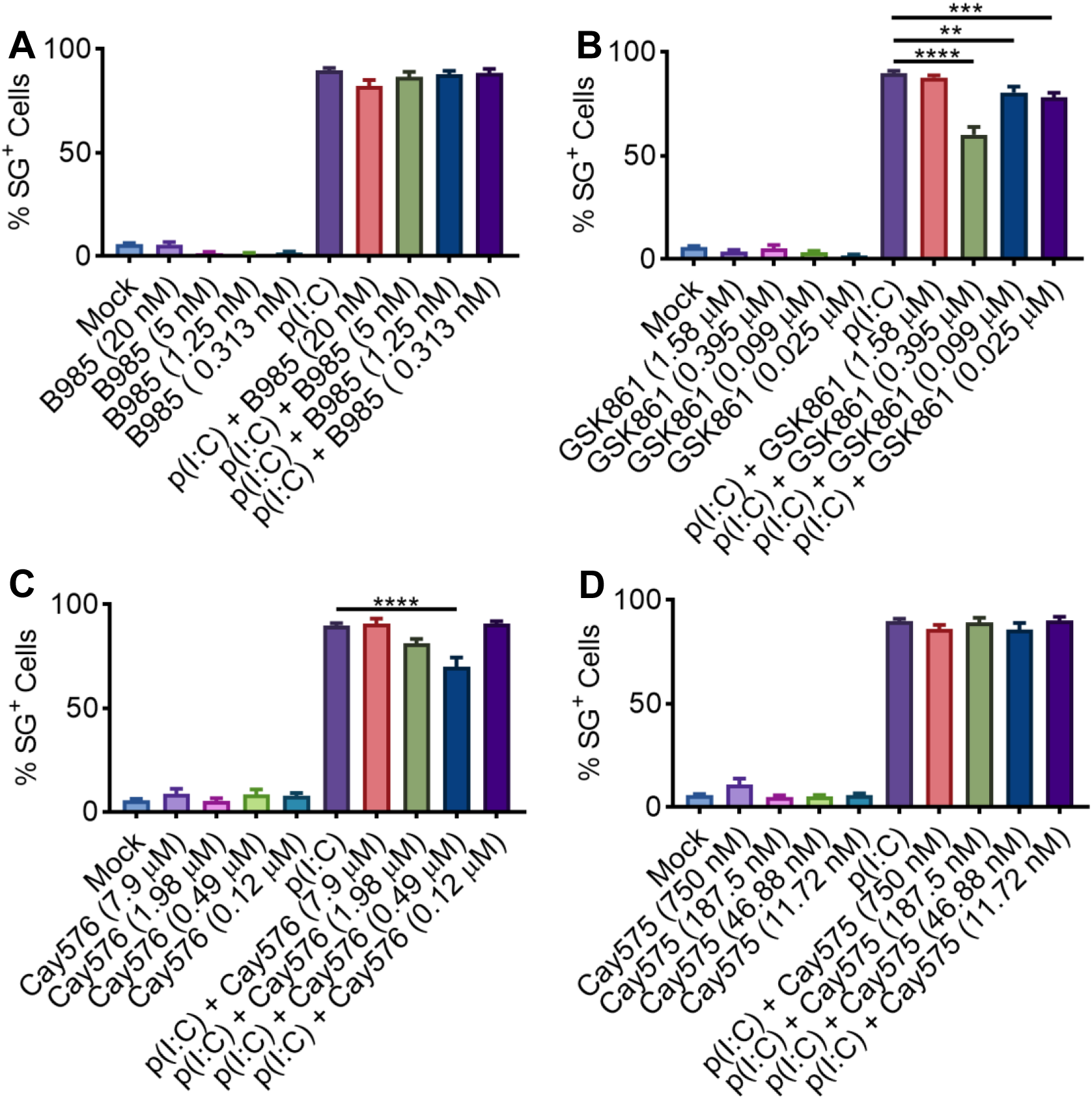
Two of the three TBK1 inhibitors destabilize SGs. **A**) Quantification of SG^+^ cells after BAY-985 (B985) treatment. BAY-985 is an inhibitor of TBK1 and IKKε kinases. **B**) Quantification of SG^+^ cells after GSK8612 (GSK861) treatment. GSK8612 is an inhibitor of TBK1 kinase. **C**) Quantification of SG^+^ cells after Cay10576 (Cay576) treatment. Cay10576 is an inhibitor of TBK1, IKKβ and PLK1 kinases. **D**) Quantification of SG^+^ cells after Cay10575 (Cay575) treatment. Cay10575 is an inhibitor of IKKβ and PLK1 kinases. One-way ANOVA was used to determine the statistical significance. \*\**p*-value<=0.01, \*\*\**p*-value<=0.001, and \*\*\*\**p*-value<=0.0001.

RIPK3, a multi-functional protein kinase, is a central component of the necroptosis RCD pathway. It can also be activated downstream of TLR signaling. Interestingly, our screen identified RIPK3 as a critical player in SG destabilization. Inhibition of RIPK3 with specific small molecule inhibitors, GSK872 and HS-1371, led to significant destabilization of SGs **(Figure 4A-B**). A recent study has reported that SGs act as a platform for activating necroptosis in interferon primed cells^42^. Our finding in unprimed cells highlights another potential intersection point between SGs and necroptotic pathways in immune response modulation.

**Figure 4:**
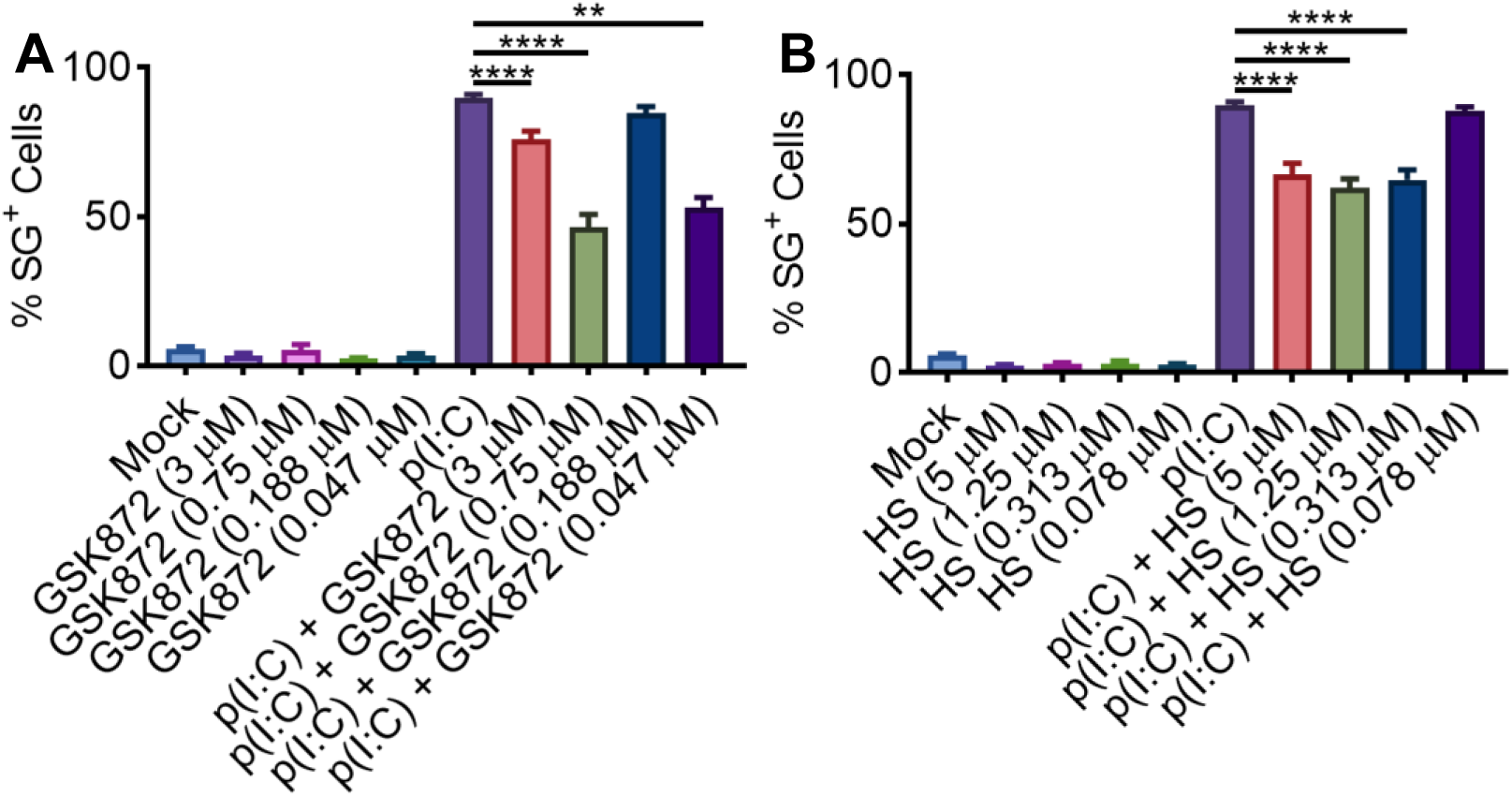
Inhibition of RIPK3 destabilizes SGs. **A**) Quantification of SG^+^ cells after GSK872 treatment. GSK872 is an inhibitor of RIPK3 kinase. **B**) Quantification of SG^+^ cells after HS-1371 (HS) treatment. HS-1371 is an inhibitor of RIPK3 kinase. One-way ANOVA was used to determine the statistical significance. \*\**p*-value<=0.01 and \*\*\*\**p*-value<=0.0001.

Beyond the canonical TLR signaling pathway, AKT signaling pathway can also be activated downstream of TLRs. AKT is a central regulator of cell growth, metabolism, and survival. To test the effect of AKT kinase activity on SG stability, we used two small molecule inhibitors – AKT inhibitor XI and Cay10567. Treatment with AKT inhibitor XI destabilized SGs while Cay10567 had no significant effect (**Figure 5A-B**). The role of AKT in SG regulation may be linked to protein synthesis and cellular metabolism, which are critical during cellular stress.

**Figure 5:**
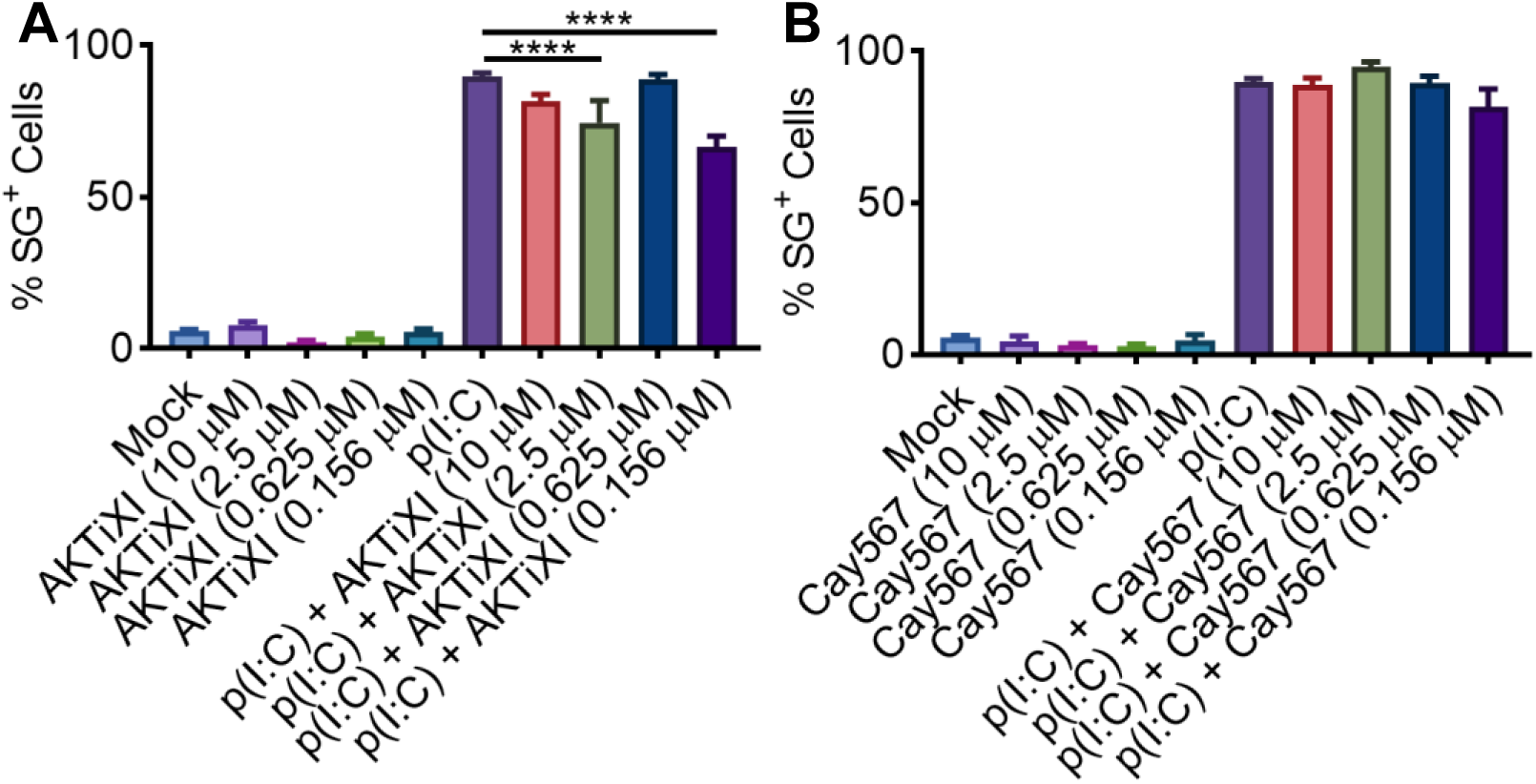
AKT inhibitors have a differential effect on SG stability. **A**) Quantification of SG^+^ cells after AKT inhibitor XI (AKTiXI) treatment. AKT inhibitor XI is an inhibitor of AKT kinase. **B**) Quantification of SG^+^ cells after Cay10567 (Cay567) treatment. Cay10567 is an inhibitor of AKT kinase. One-way ANOVA was used to determine the statistical significance. \*\*\*\**p*-value<=0.0001.

### MAP kinases inhibitors influence SG stability

MAP kinases are the evolutionary highly conserved signaling pathways that are involved in several aspects of immune responses. MAP kinases signaling is activated by various stimuli, including TLRs. There are 3 MAP kinases – c-Jun N-terminal kinase (JNK), extracellular signal-Regulated Kinase (ERK) and P38. In our study, we tested the effect of inhibitors targeting JNK1/2/3 and ERK1/2 kinases on SG stability.

Interestingly, the inhibition of JNK1/2/3 kinases by JNK inhibitor VII or JNK inhibitor VIII destabilized SGs. (**Figure 6A-B**). For ERK1/2, our data shows that inhibition of ERK1/2 kinases by two specific inhibitors, LY3214996 and FR180204, results in different effects on SG dynamics. Treatment with LY3214996 significantly reduced the number of SG-positive cells **(Figure 6C)**, indicating that ERK1/2 inhibition decreases SG stability. On the other hand, treatment with FR180204, another ERK1/2 inhibitor, led to an increase in the number of SG-positive cells indicating SG stabilization **(Figure 6D)**. In our previous study, we had observed that ERK1/2 activity promotes SG assembly in BMDMs in response to ER-stress^9^. However, the contrasting effects of these two inhibitors raise the possibility that ERK1/2 may have a pleiotropic effect on SG assembly and stability. This differential response could be due to selective targeting of specific downstream substrates of ERK1/2 kinases, such as SRSF1 and SRSF3, which are involved in RNA metabolism and SG formation. Another possibility is that one of the two inhibitors also reduces the activity of another unknown kinase involved in SG biology. Taken together our results have uncovered a key link between the MAP kinase signaling pathway and SG regulation.

**Figure 6:**
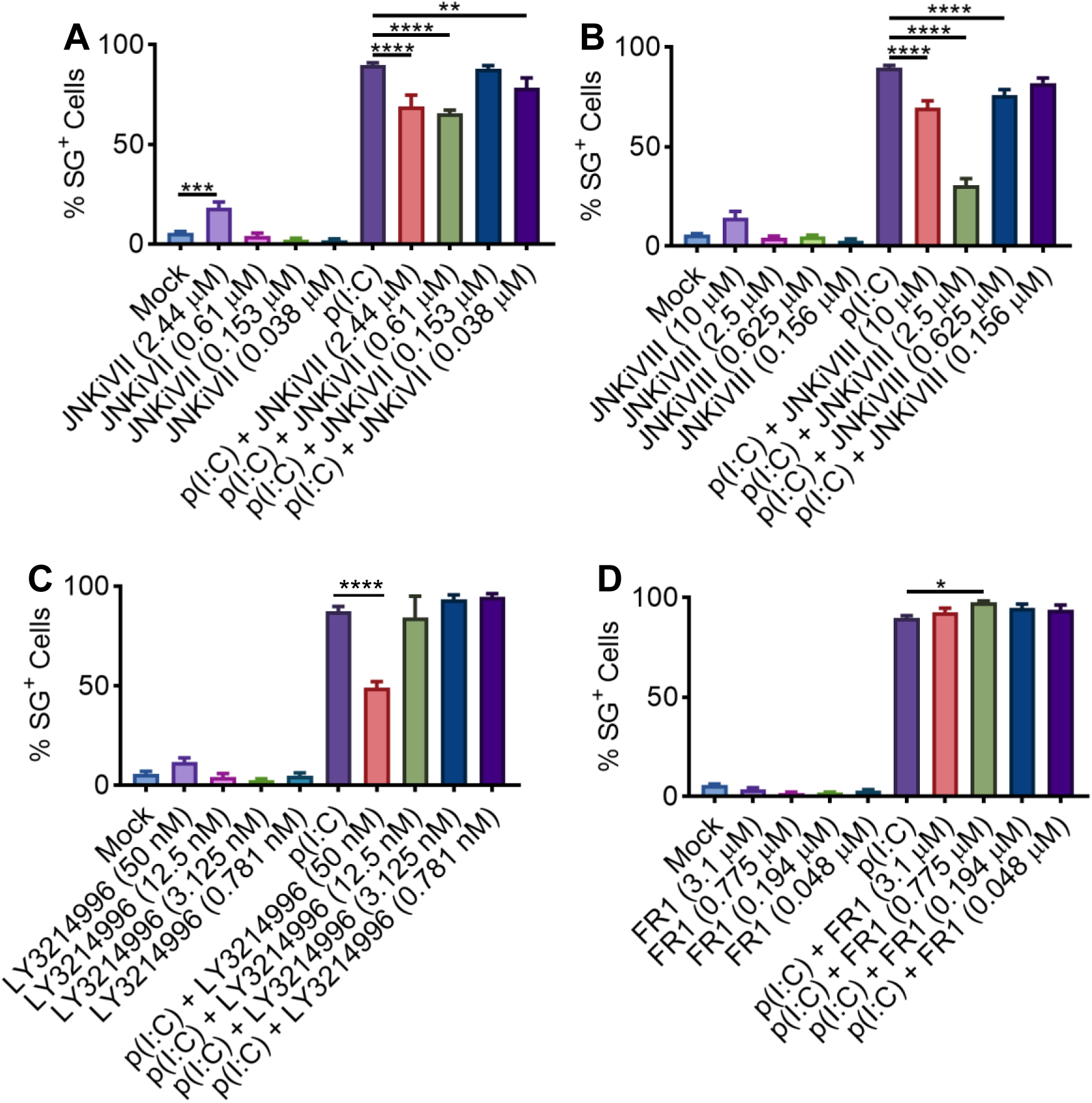
MAP kinases differentially modulate SG stability. **A**) Quantification of SG^+^ cells after JNK inhibitor VII (JNKiVII) treatment. JNK inhibitor VII is an inhibitor of JNK1, JNK2 and JNK3 kinases. **B**) Quantification of SG^+^ cells after JNK inhibitor VIII (JNKiVIII) treatment. JNK inhibitor VIII is an inhibitor of JNK1, JNK2 and JNK3 kinases. **C**) Quantification of SG^+^ cells after LY3214996 treatment. LY3214996 is an inhibitor of ERK1/2 kinases. **D**) Quantification of SG^+^ cells after FR180204 (FR1) treatment. FR180204 is an inhibitor of ERK1/2 kinases. One-way ANOVA was used to determine the statistical significance. \**p*-value<=0.05, \*\**p*-value<=0.01 and \*\*\*\**p*-value<=0.0001.

### PERK increases activity of SGs assembled due to PKR activation

PERK is a key sensor of endoplasmic reticulum (ER) stress and plays an important role in the unfolded protein response (UPR). PERK activation leads to phosphorylation of eIF2α, which inhibits global protein synthesis but facilitates the translation of stress-related proteins. Interestingly, TLR stimulation has also been shown to activate ER stress signaling, albeit through a different stress sensor named XBP1^43^. Intriguingly, the inhibition of PERK kinase by GSK2606414 or GSK2656157 destabilized SGs formed in response to dsRNA sensing (**Figure 7A-B**). The destabilization of SGs upon PERK inhibition highlights its critical role in maintaining SG integrity, especially under stress conditions such as those induced by viral infection or ER stress. This also suggests that stress sensing kinases can reinforce signaling induced by others in the stress sensing kinase network to modulate SG assembly and stability.

**Figure 7:**
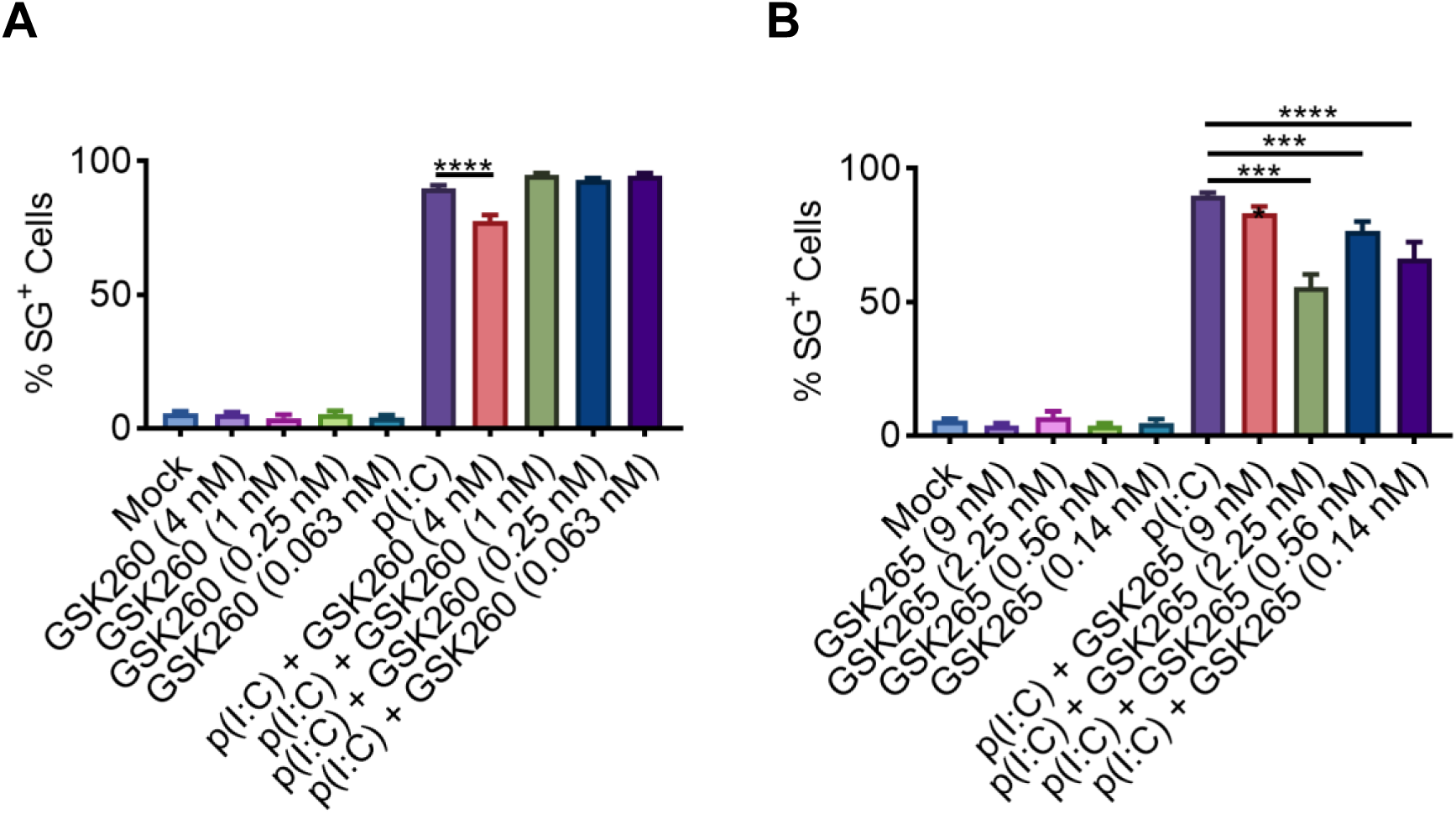
PERK reinforces PKR triggered stress signaling. **A**) Quantification of SG^+^ cells after GSK2606414 (GSK260) treatment. GSK2606414 is an inhibitor of PERK kinase. **B**) Quantification of SG^+^ cells after GSK2656157 (GSK265) treatment. GSK2656157 is an inhibitor of PERK kinase. One-way ANOVA was used to determine the statistical significance. \*\*\**p*-value<=0.001 and \*\*\*\**p*-value<=0.0001.

### Context-dependent regulation of SG by NF-κB Signaling

The NF-κB signaling pathway is a crucial regulator of inflammation, immune responses, and cellular stress that is also activated by TLR signaling. Strikingly, our data demonstrates that the inhibition of NF-κB signaling pathway has differential effects on SG dynamics. Inhibition of the NF-κB pathway with different inhibitors revealed both spontaneous induction and destabilization of SGs. For instance, PPM18, an NF-κB inhibitor, induced spontaneous SG formation while also promoting the disassembly of SGs triggered by dsRNA sensing **(Figure 8A).** A similar effect was observed with BAY-11-7082, another NF-κB inhibitor, which induced spontaneous SGs but destabilized those formed in response to poly(I:C) transfection **(Figure 8B).** Interestingly, RO-106-9920, another inhibitor targeting the NF-κB pathway, did not induce spontaneous SGs but destabilized pre-formed SGs **(Figure 8C).** In contrast, other inhibitors such as BMS-345541 and QNZ showed no measurable effect on SG assembly or disassembly **(Figure 8D-E).**

**Figure 8:**
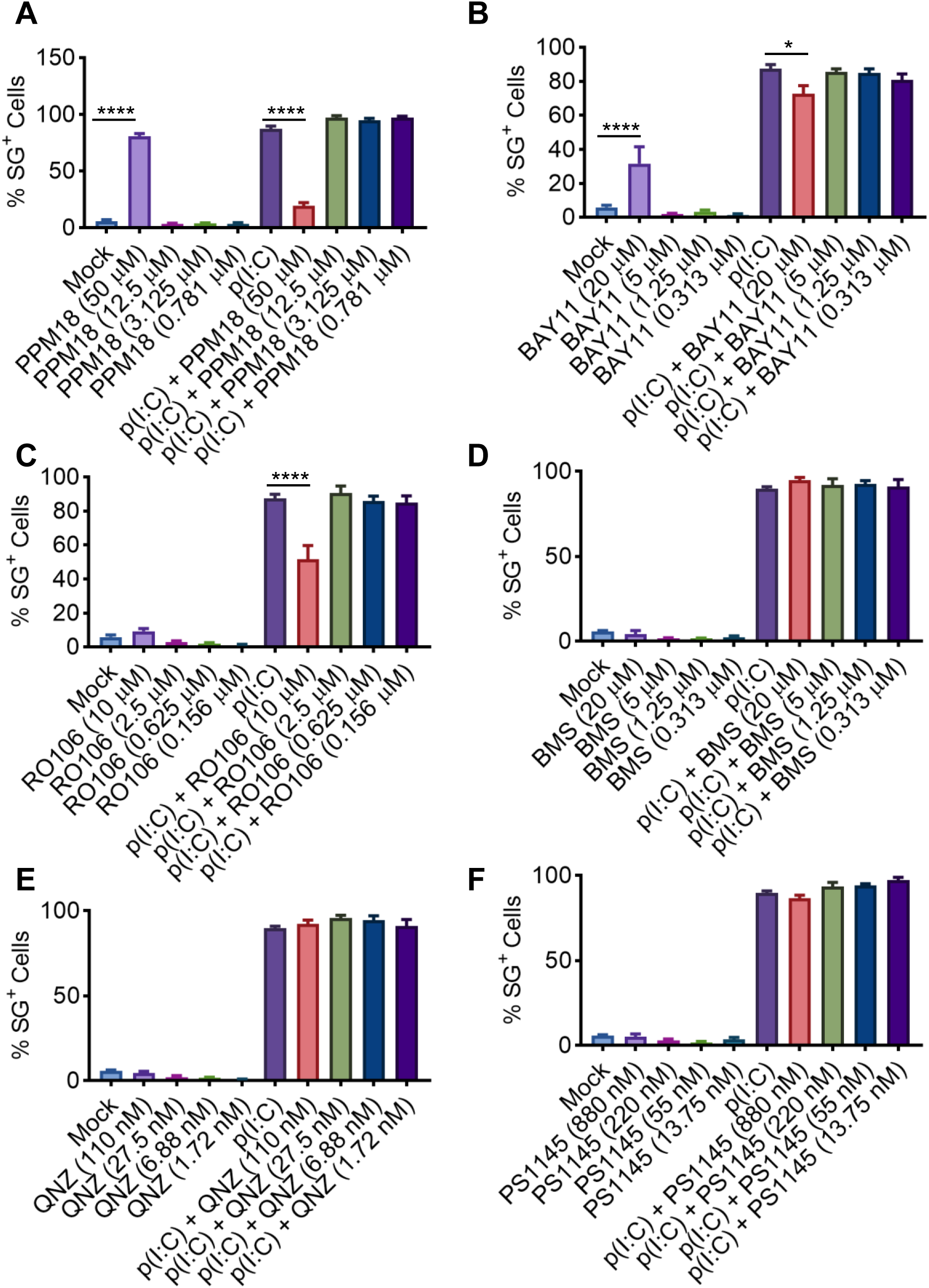
IKK1/2 inhibitors differentially modulate SG stability. **A**) Quantification of SG^+^ cells after PPM18 treatment. PPM18 is an inhibitor of the NF-κB signaling pathway. **B**) Quantification of SG^+^ cells after BAY-11-7082 (BAY11) treatment. BAY-11-7082 is an inhibitor of the NF-κB signaling pathway. **C**) Quantification of SG^+^ cells after RO-106-9920 (RO106) treatment. RO-106-9920 is an inhibitor of the NF-κB signaling pathway. **D**) Quantification of SG^+^ cells after BMS-345541 HCl (BMS) treatment. BMS-345541 HCl is an inhibitor of the NF-κB signaling pathway. **E)** Quantification of SG^+^ cells after QNZ treatment. QNZ is an inhibitor of the NF-κB signaling pathway. **F**) Quantification of SG^+^ cells after PS1145 treatment. PS1145 is an inhibitor of IKKβ kinase. One-way ANOVA was used to determine the statistical significance. \**p*-value<=0.05 and \*\*\*\**p*-value<=0.0001.

These contrasting outcomes suggest that these NF-κB inhibitors could be modifying IKK complex substrate specificity to target distinct downstream components of the pathway, possibly reflecting differential inhibition of IKKα and IKKβ kinases. Supporting this hypothesis, we found that treatment with Cay10575 (**Figure 3D**) or PS1145 (**Figure 8F**) which inhibit the IKKβ kinase does not affect SG stability. This suggests that the IKKα kinase, rather than IKKβ, may be playing a more prominent role in SG biology. In summary, NF-κB signaling appears to play a complex role in SG regulation.

### PPM18 induces eIF2α phosphorylation and forms SGs in a dose dependent manner

Surprisingly, our data shows that two of the NF-κB inhibitors, PPM18 and Bay-11-7082, induced spontaneous SG formation even in the absence of external stress. Interestingly, BAY-11-7082 had previously been reported to inhibit IAV replication^44^. To investigate the underlying mechanisms of SG formation and its potential antiviral applications, we focused on PPM18, as it robustly induces SGs in a majority of cells. To determine if PPM18 induces SGs in a dose dependent manner, we treated A549 cells with increasing concentrations of the compound. Our results show a clear dose-dependent increase in the number of SG positive cells, indicating that SG formation is linked to PPM18 treatment. (**Figure 9A-B**). SG assembly can be influenced by modulating polysome stability. While stabilization of polysomes (e.g., through translational elongation inhibitor) prevents SG formation, inhibition of translation initiation promotes it. To assess if PPM18 induced SGs are similarly affected by polysome stabilization, we treated A549 cells with either cycloheximide (elongation inhibitor) or puromycin (initiation inhibitor). Cycloheximide treatment strongly inhibited SGs induced by PPM18 whereas puromycin did not have any effect, suggesting that PPM18 engages SG assembly pathway dependent upon translation initiation inhibition (**Figure 9C-D**). Next, to determine if PPM18 induces engages eIF2α-dependent SG formation, we measured eIF2α phosphorylation using western blot analysis. Phosphorylation of eIF2α is the initiator event in the canonical SG assembly pathway that relies on translation initiation inhibition due to disruption of eIF2B activity^45,46^. PPM18 treatment indeed induces robust eIF2α phosphorylation indicating that the compound triggers the canonical SG assembly pathway (**Figure 10A**). There are four eIF2α kinases that sense different types of stress and phosphorylate eIF2α to activate the integrated stress response (ISR) signaling pathway. ISR activation is upstream of SG assembly and the three ISR kinases (EIF2AK1, EIF2AK2 and EIF2AK3) play a central role in inducing SGs as well as EIF4AK4 which can also phosphorylate eIF2α^47,48^. To identify which eIF2α kinase was responsible for initiating SG assembly, we generated knockout A549 cells and validated the three SG inducing ISR kinases using western blot analyses (**Figure 10B-D**). Our data shows a marked decrease in SG positive cells in absence of EIF2AK3 (**Figure 10E-F**). Loss of EIF2AK1 also results in a lesser number of SG positive cells, however this decrease is significantly weaker in the case of EIF2AK3 (**Figure 10E-F**). Since EIF2AK3 senses endoplasmic reticulum stress (ER-stress) and EIF2AK1 is activated in response to a variety of stresses including oxidative stress, our data could be indicating that PPM18 treatment induces both ER and oxidative stresses. However, our observation with PERK inhibitors (**Figure 7A-B**) indicates that stress sensing kinases can reinforce each other’s signaling within the ISR network. The nearly complete loss of SG formation in *EIF2AK3*^-/-^ cells, combined with partial loss in *EIF2AK1*^-/-^ cells (**Figure 10E-F**), raises an alternate possibility that HRI amplifies PERK mediated stress signaling rather than acting independently. Taken together, our data shows that PPM18 induces SGs by inducing eIF2α phosphorylation by activating PERK (EIF2AK3) and/or HRI (EIF2AK1).

**Figure 9:**
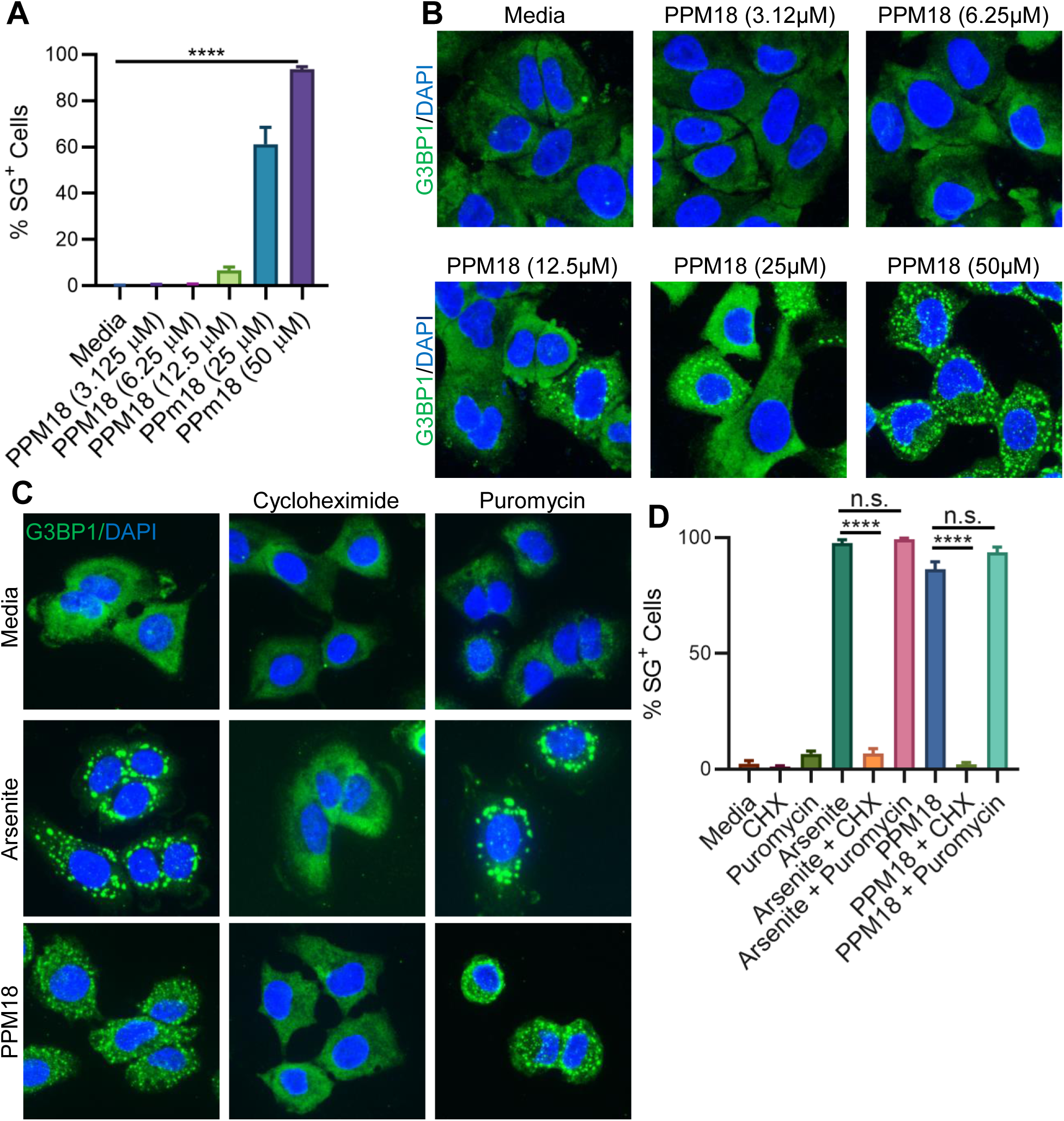
PPM18 induced spontaneous SG are dose dependent and inhibited by polysome stabilization. **A**) Quantification of SG^+^ cells after PPM18 treatment. **B**) Representative images of cells from the treatment groups quantified in panel **A**. **C**) Representative images of SG^+^ cells upon PPM18 treatment following inhibition of translation elongation (cycloheximide) or translation initiation (puromycin). Sodium arsenite treatment was used as positive control. **D**) Quantification of cells from the treatment groups shown in panel **A**. CHX is cycloheximide. G3BP1 was used as the SG marker. DAPI was used to visualize nuclei. One-way ANOVA was used to determine the statistical significance. \*\*\*\**p*-value<=0.0001 n.s.: not significant.

**Figure 10:**
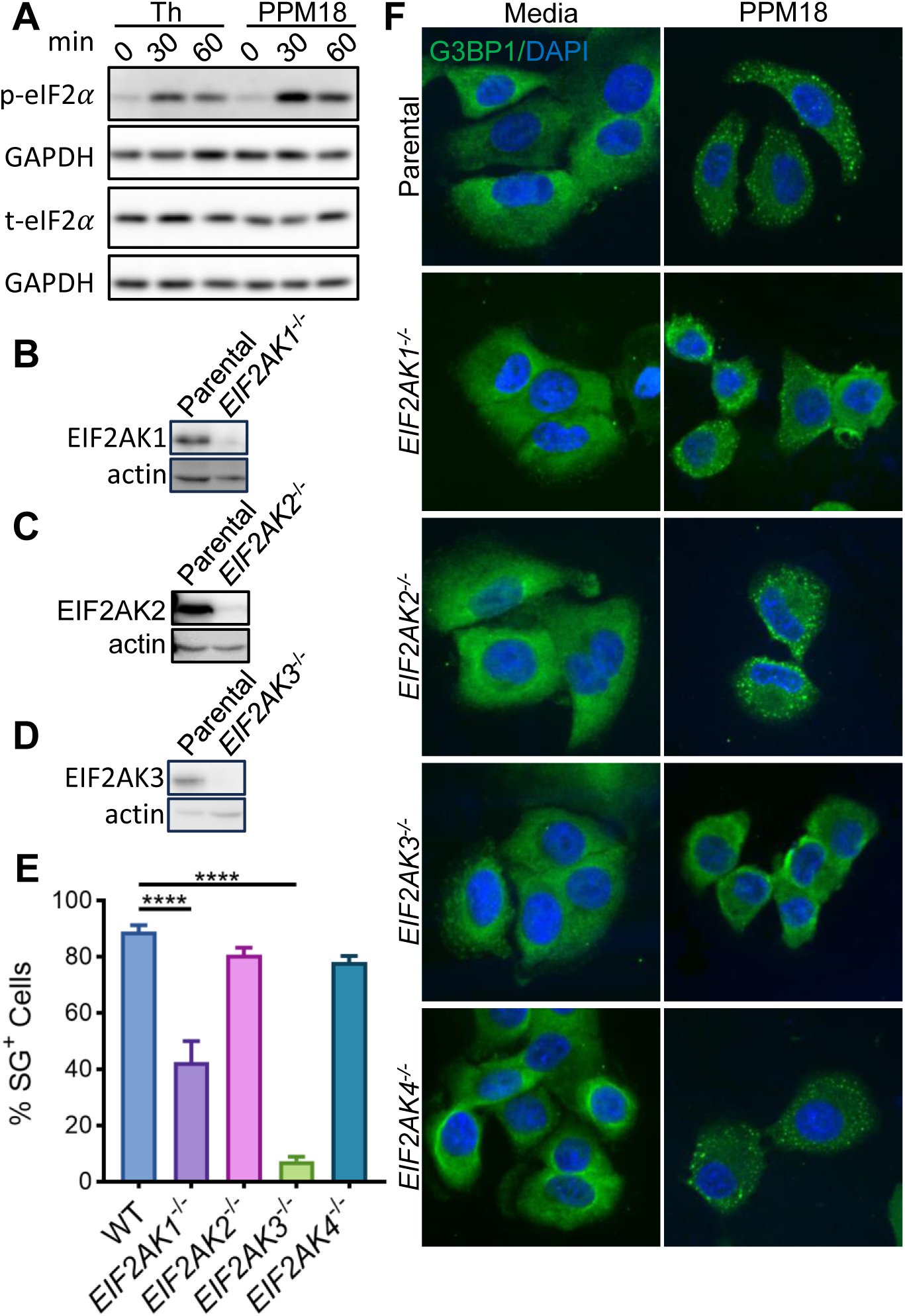
PPM18 induces SG by activating EIF2AK1 and EIF2AK3. **A**) Western blot analysis of eIF2α phosphorylation in response to PPM18 treatment. **B-D**) Validation of gene knockouts in cells. *EIFAK1*, *EIFAK2*, and *EIFAK3* were knocked out in A549 cells. **E**) Quantification of SG^+^ cells after PPM18 treatment. Untransduced parental A549 cells were used as WT control. **F**) Representative images of cells from the treatment group quantified in panel **E**. G3BP1 was used as the SG marker. DAPI was used to visualize nuclei. One-way ANOVA was used to determine the statistical significance. \*\*\*\**p*-value<=0.0001.

### PPM18 inhibits IAV replication

A major goal of this project was to identify small molecules that can be used as a host directed antiviral drug. To test the antiviral potential of PPM18, we treated A549 cells with the compound after allowing IAV to adsorb for 2 hours. Excitingly, PPM18 robustly inhibited IAV replication 9 hours post infection, and the viral replication inhibition was effective even after 24 hours of infection (**Figure 11A-B**). Since IAV replication occurs in the nucleus, while SG assembles in the cytoplasm, we investigated if PPM18 inhibits IAV replication by changing the nucleocytoplasmic distribution of viral RNPs (vRNPs) using immunofluorescence confocal microscopy. Our results show that IAV signal remained predominantly nuclear in infected cells treated with PPM18, indicating that the antiviral effect is not due to altered vRNP redistribution (**Figure 11C**). This indicates that viral replication inhibition by PPM18 is through a distinct mechanism unrelated to vRNP nucleocytoplasmic transport.

**Figure 11:**
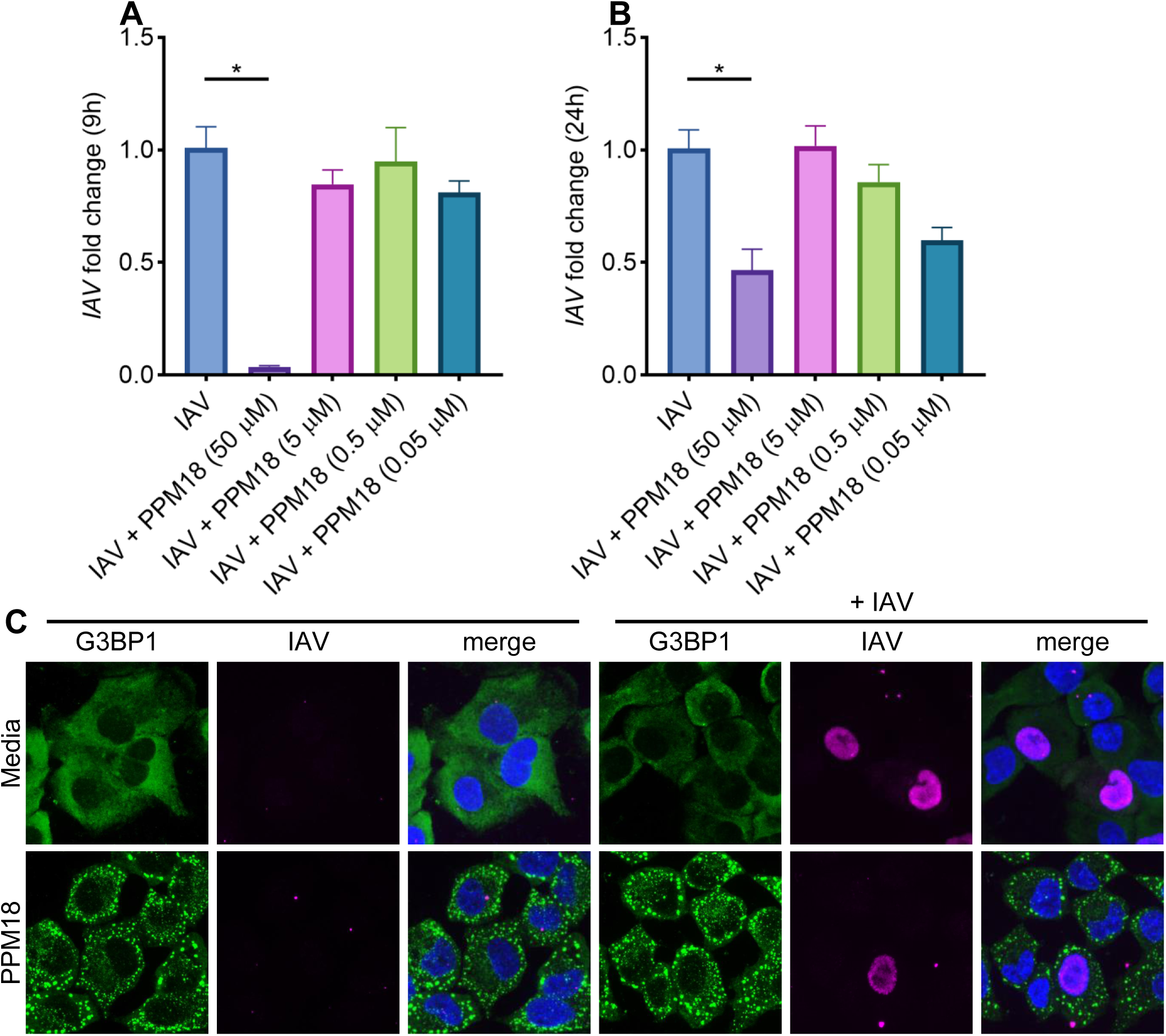
PPM18 inhibits IAV replication without affecting sub-cellular localization of IAV vRNPs. **A**) qPCR measurement of viral RNA 9 hours post infection as a marker for viral replication. PPM18 was added to cells 2 hours after infection to ensure we are measuring effect of PPM18 on viral replication and not infection efficiency. Cells were infected at multiplicity of infection (M.O.I.) of 0.1. **B**) qPCR measurement of viral RNA 24 hours post infection as a marker for viral replication. PPM18 was added to cells 2 hours after infection as in panel **A**. Cells were infected at multiplicity of infection (M.O.I.) of 0.1. **C**) Visualization of sub-cellular localization of IAV viral ribonucleoprotein complexes (vRNPs) after PPM18 treatment. Cells were infected with IAV at M.O.I. of 1. G3BP1 was used as the SG marker. DAPI was used to visualize nuclei. vRNPs were visualized using a goat polyclonal antibody raised against IAV. One-way ANOVA was used to determine the statistical significance. \**p*-value<=0.05.

### PPM18 induces caspase independent regulated cell death at antiviral concentrations

We observed that PPM18 treatment was leading to extensive cell rounding probably due to simultaneous activation of two stress sensing kinases EIF2AK1 and EIF2AK3. Prolonged stress signaling can induce RCD in cells. To test if PPM18 was inducing RCD, we performed live-cell imaging in presence of cell death reporter dye propidium iodide. We used extrinsic apoptosis induced by TNF + cycloheximide treatment as a positive control (**Figure 12A,C**). PPM18 induced RCD in A549 cells at the concentration (50 µM) that effectively inhibited IAV replication (**Figure 12B,C**). Although, PPM18 cytotoxicity greatly reduces its translational potential as an antiviral drug, potential solution could be to co-administer it with an RCD inhibitor that preserves its antiviral effect while preventing cell death. Therefore, we tested if a pan-caspase inhibitor zVAD-fmk can interfere with PPM18 induced RCD. However, our data indicates that zVAD-fmk failed to inhibit PPM18 induced RCD in A549 cells (**Figure 12B,C**), despite completely inhibiting extrinsic apoptosis induced by TNF + cycloheximide treatment (**Figure 12A,C**). In summary, PPM18 induces RCD in A549 cells through a caspase-independent pathway.

**Figure 12:**
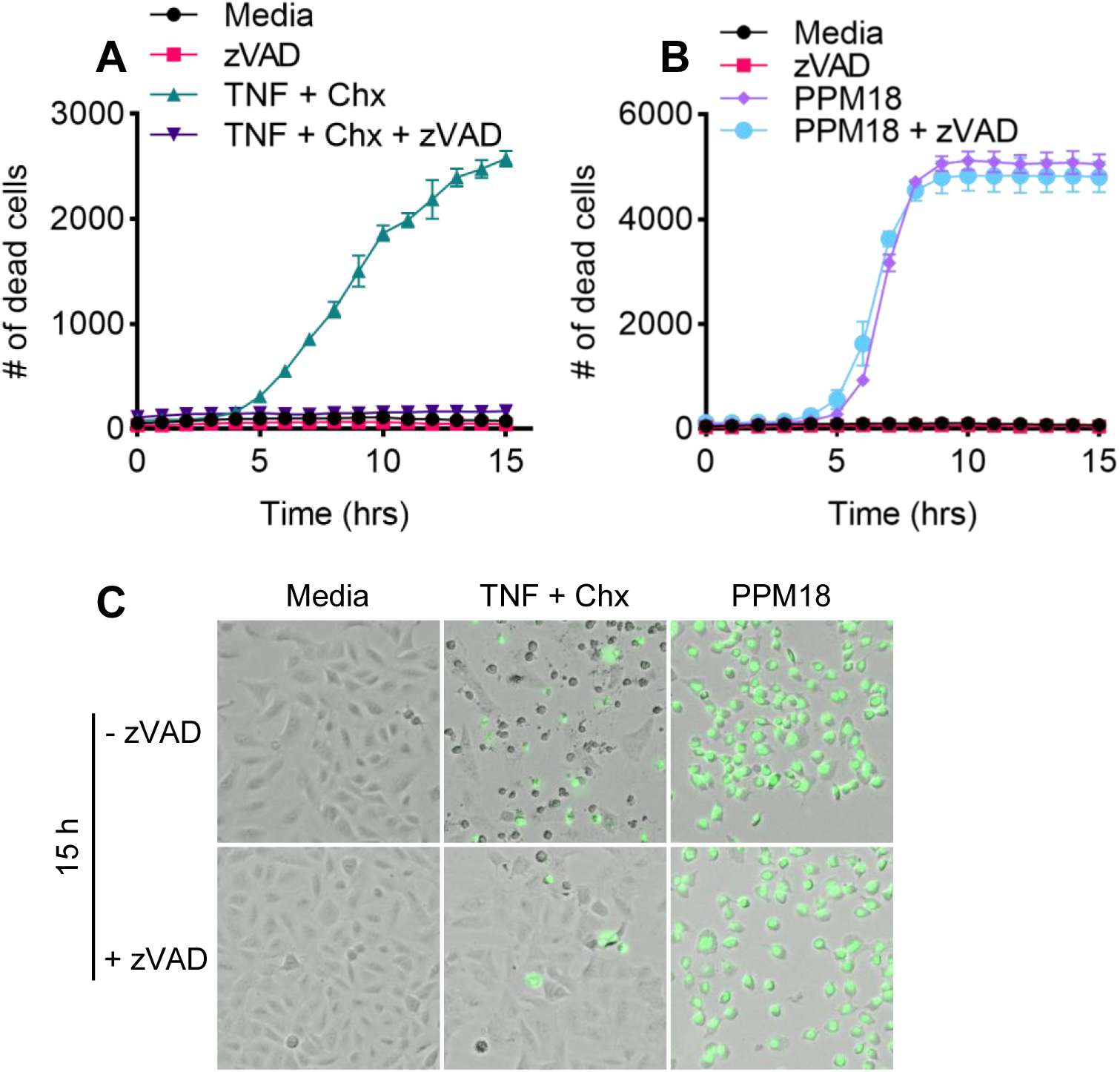
PPM18 induced RCD is not affected by pan-caspase inhibitor zVAD-fmk. **A**) Propidium iodide staining based measurement of RCD in A549 cells treated with TNF + cycloheximide (Chx). Pan-caspase inhibitor zVAD-fmk-oMe was used to inhibit RCD induced by TNF + cycloheximide treatment. **B**) Propidium iodide staining based measurement of RCD in A549 cells treated with PPM18. Pan-caspase inhibitor zVAD-fmk-oMe was used to inhibit caspases to determine if RCD induced by PPM18 was dependent on caspase activation. **C**) Representative images showing live-dead cells quantified in panels **A** and **B**. Dead cells are pseudo-colored green based on propidium iodide staining.

### PPM18 induces robust *TNF* and *IFNB1* expression

SGs have been reported to have conflicting roles in type I interferon signaling, with some studies suggesting enhancement of type I interferon signaling while others suggest suppression^12,16,27,28^. PPM18 has previously been reported to inhibit inflammatory signaling in bone marrow derived macrophages in response to lipopolysaccharide stimulation^49^. Since IAV infection induces a complex cytokine response including a robust type I interferon signaling activation, we examined weather PPM18 treatment alters pro-inflammatory gene expression. qPCR analysis revealed that PPM18 treatment significantly increased mRNA abundances of *IFNB* and *TNF* 9 hours post infection (**Figure 13A,C**). This effect of PPM18 on gene expression of the pro-inflammatory cytokines was even stronger at later timepoints, suggesting that PPM18 enhances antiviral and proinflammatory innate immune signaling (**Figure 13B,D**). Our results in combination with the published study indicate a differential effect of PPM18 on inflammatory gene expression based on cell types and/or PRRs that are activated.

**Figure 13:**
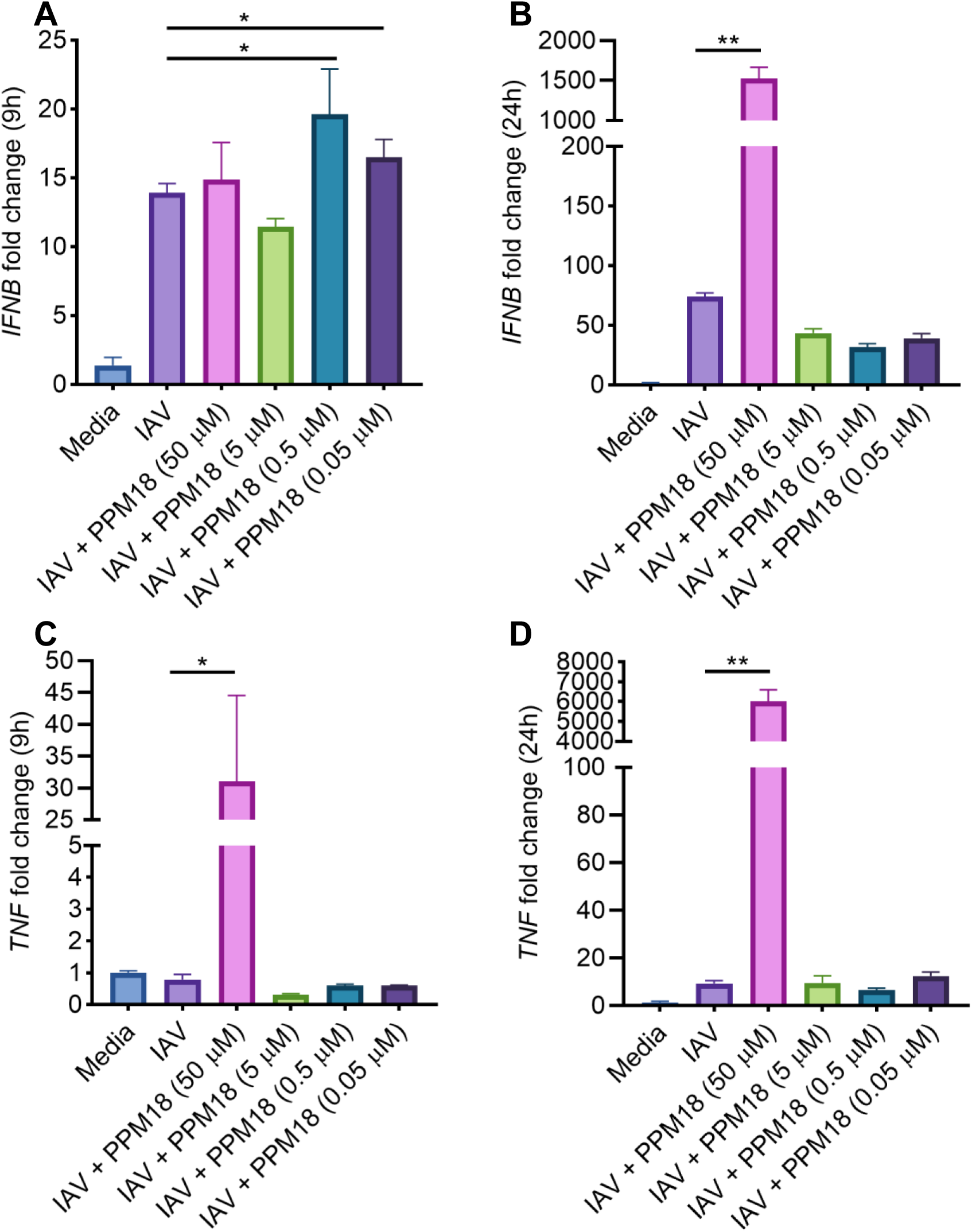
PPM18 induces robust TNF and IFNB expression. **A**) qRT-PCR measurement of *IFNB* expression in PPM18 treated IAV infected cells 9 hours post infection. PPM18 was added to cells 2 hours post infection. **B**) qRT-PCR measurement of *IFNB* expression in PPM18 treated IAV infected cells 24 hours post infection. PPM18 was added to cells 2 hours post infection. **C**) qRT-PCR measurement of *TNF* expression in PPM18 treated IAV infected cells 9 hours post infection. PPM18 was added to cells 2 hours post infection. **D**) qRT-PCR measurement of *TNF* expression in PPM18 treated IAV infected cells 24 hours post infection. PPM18 was added to cells 2 hours post infection. One-way ANOVA was used to determine the statistical significance. \**p*-value<=0.05 and \*\**p*-value<=0.01.

### PPM18 mediated viral replication inhibition and RCD induction are independent of SGs

SGs have been reported to inhibit IAV replication in several studies. Since PPM18 effectively inhibits IAV replication at concentration that also induces SGs, we sought to determine if SGs play a role in this antiviral effect. To test the role of SGs in PPM18 mediated IAV replication inhibition, we generated *G3BP1*^-/-^ A549 cells (**Figure 14A**). We prepared monoclonal cultures of the knockout cells and used three independent clones in our experiments (**Figure 14A**). Loss of G3BP1 abrogated SG assembly induced by PPM18 treatment indicating that presence of G3BP1 was necessary for PPM18 triggered SG assembly (**Figure 14B-C**). We then tested if PPM18 was able to inhibit IAV replication in *G3BP1*^-/-^ cells. Strikingly, PPM18 still inhibited IAV replication in all the three clones that we tested (**Figure 15A-C**), suggesting that SGs are not required for antiviral activity. Interestingly, we observed a trend towards increased IAV replication in untreated *G3BP1*^-/-^ cells compared to the parental WT A549 cells. This indicates that G3BP1 might be playing a role in restricting IAV replication at basal level. However, since IAV actively inhibits SG formation via IAV-NS1 protein mediated sequestration of host DDX3X protein, SGs are not detected in IAV infected cells. Therefore, it is unlikely that G3BP1 restricts viral replication through its role as a SG scaffold protein. The exact mechanism by which G3BP1 interferes with viral replication is a subject for future studies. Although previous studies had reported that SGs protect against RCD, a more recent study had reported that they can also act as a platform for activating necroptosis^4,5,42,50^. Since PPM18 mediated RCD is caspase independent, and it is a robust inducer of SGs, we tested role of SGs in PPM18 mediated RCD using *G3BP1*^-/-^ cells. PPM18 still induced RCD in *G3BP1*^-/-^ cells with the same efficiency as the parental A549 cells indicating SGs do not play a role in this process (**Figure 15D-I**). In summary, our results show that PPM18 induces SGs by activating ISR but inhibits IAV replication through a mechanism that does not depend on SGs.

**Figure 14:**
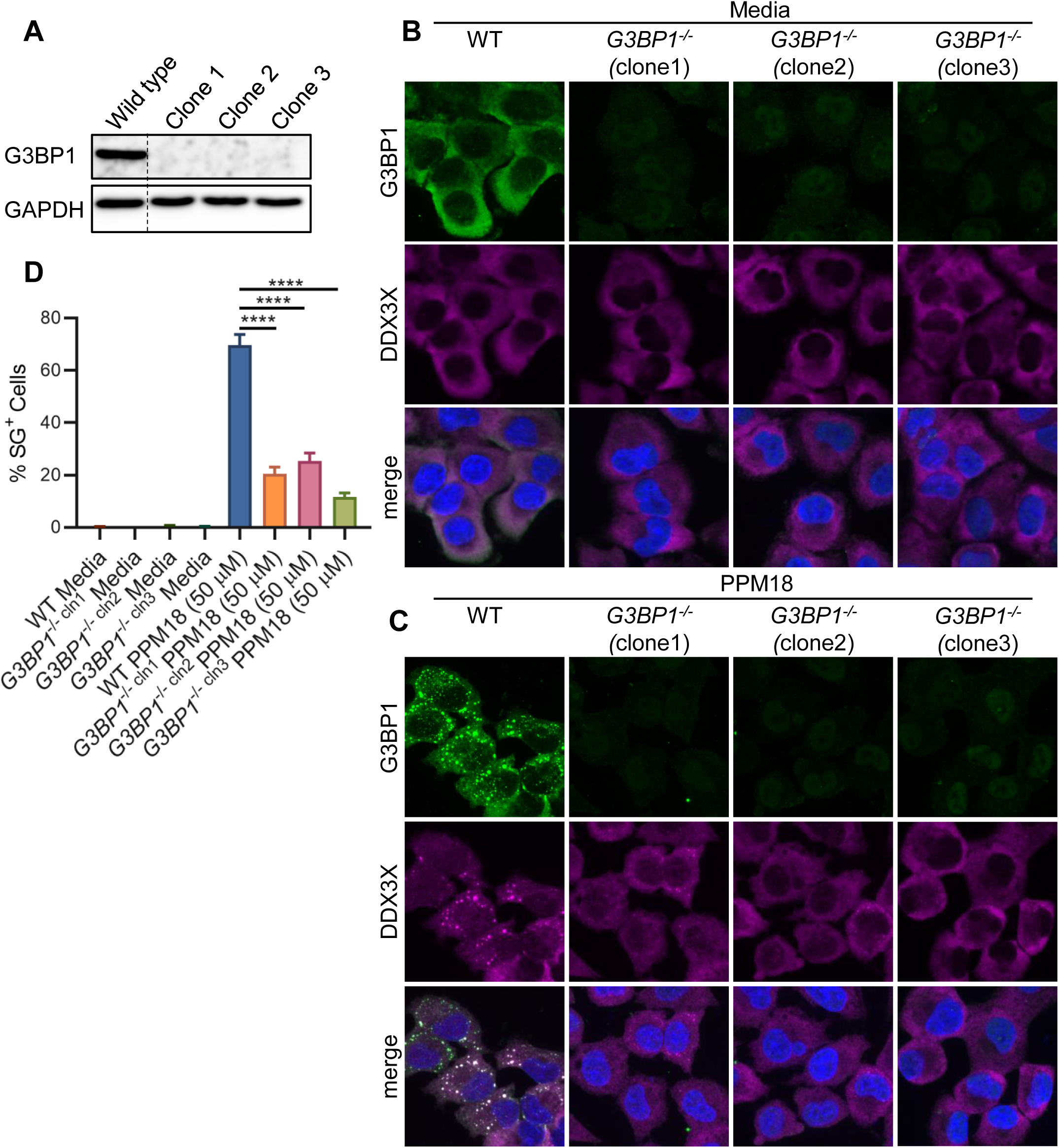
PPM18 mediated SG assembly is dependent on G3BP1. **A**) Western blot validation of gene knockouts for the three *G3BP1*^-/-^ A549 cells clones. **B**) Immunofluorescence confocal imaging of mock treated WT and *G3BP1*^-/-^ A549 cells to visualize G3BP1 expression and SGs. **C**) Immunofluorescence confocal imaging of PPM18 treated WT and *G3BP1*^-/-^ A549 cells to visualize G3BP1 expression and SGs. **D**) Quantification of SG^+^ WT and *G3BP1*^-/-^ A549 cells upon PPM18 treatment. One-way ANOVA was used to determine the statistical significance. \*\*\*\**p*-value<=0.0001. G3BP1 was immunostained with AlexaFluor488 secondary antibody. DAPI was used to visualize nuclei. DDX3X used as the alternate SG marker.

**Figure 15:**
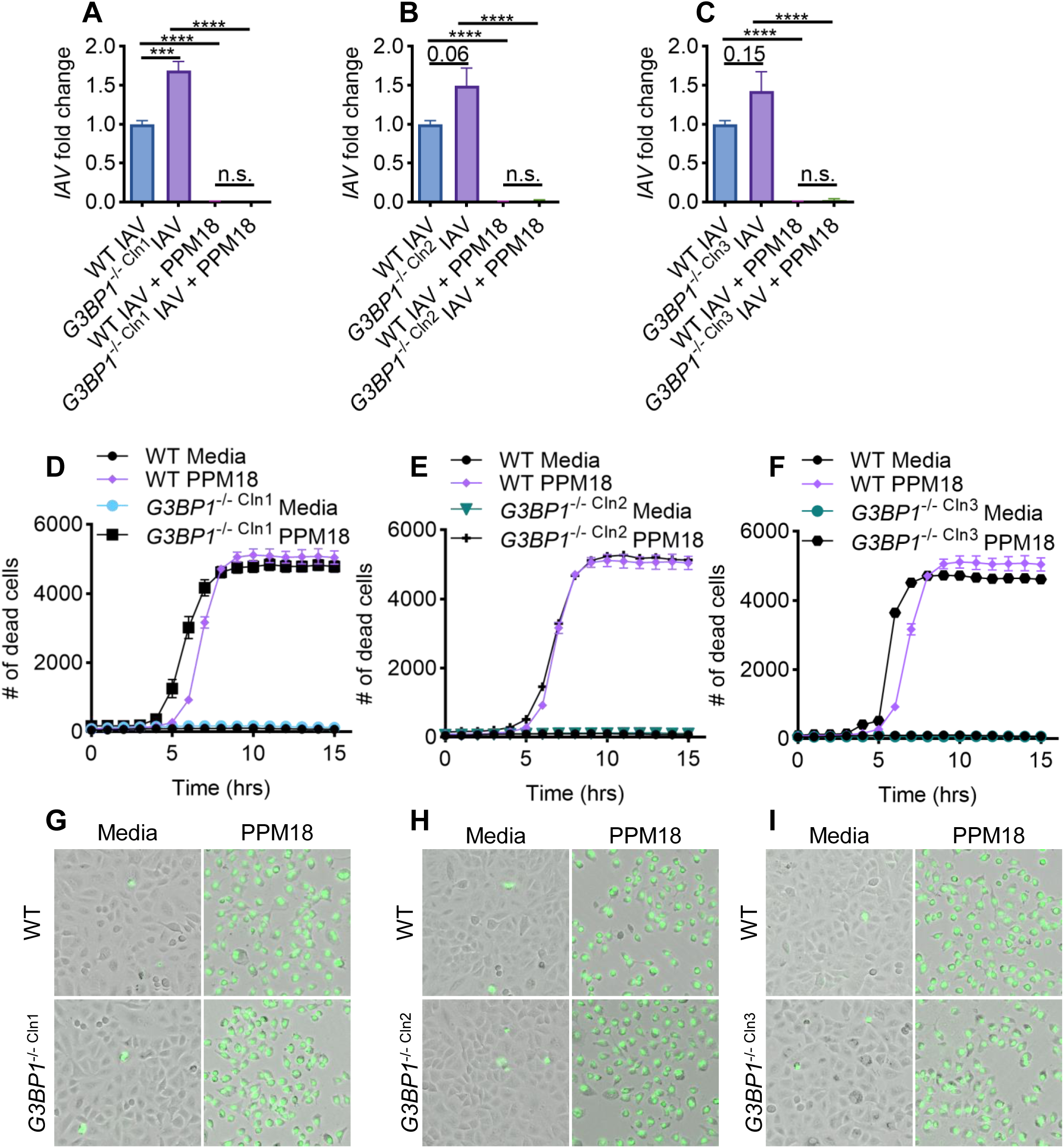
PPM18 mediated viral replication inhibition and RCD induction are independent of SG formation. **A**) qRT-PCR measurement of IAV replication in clone 1 of *G3BP1*^-/-^ cells. **B**) qRT-PCR measurement of IAV replication in clone 2 of *G3BP1*^-/-^ cells. **C**) qRT-PCR measurement of IAV replication in clone 3 of *G3BP1*^-/-^ cells. **D**) Propidium iodide staining based measurement of RCD in clone 1 of *G3BP1*^-/-^ cells. **E**) Propidium iodide staining based measurement of RCD in clone 2 of *G3BP1*^-/-^ cells. **F**) Propidium iodide staining based measurement of RCD in clone 3 of *G3BP1*^-/-^ cells. **G**) Representative images showing live-dead cells quantified in panel **D**. Dead cells are pseudo-colored green based on propidium iodide staining. **H**) Representative images showing live-dead cells quantified in panel **E**. Dead cells are pseudo-colored green based on propidium iodide staining. **I**) Representative images showing live-dead cells quantified in panel **F**. Dead cells are pseudo-colored green based on propidium iodide staining. One-way ANOVA was used to determine the statistical significance. \*\*\**p*-value<=0.001 and \*\*\*\**p*-value<=0.0001.

## Discussion

Our study has identified several small molecules that target proteins in the TLR signaling pathway and modulate stability of SGs formed in response to cytosolic dsRNA sensing, providing new insights into SG biology and its intersection with immune signaling. Notably, we found that some compounds induce spontaneous SGs, even in the absence of cytosolic dsRNA, revealing an intricate crosstalk between the TLR signaling pathway and SG dynamics that remains poorly understood.

For several kinases, one inhibitor had an effect that was different from another inhibitor for the same kinase. Several mechanisms can explain these differences. For example, the efficiency of uptake of the two inhibitors could be different, which would result in differences in their available intracellular concentrations in A549 cells. These inhibitors have different selectivity profiles, which could result in unintended targeting of unrelated kinases. In many cases, a lower dose was able to modify SG stability when a higher dose failed. This is a very interesting observation that suggests partial inhibition of a kinase might have a different effect on SG dynamics than its complete inhibition. These findings raise new questions and open new avenues for research in SG biology and its modulation by the innate immune response.

One particular intriguing finding is that that an inhibitor of ERK1/2 kinases stabilizes while another destabilizes SGs formed in response to dsRNA sensing. In a previous study, we had found that ERK1/2 also promotes assembly of SGs formed in response to ER-stress^9^. How does ERK1/2 activity promote SG assembly or stability remains unclear. One mechanism could involve phosphorylation of a SG component as several SG components, such as SRSF1, SRSF3 and DDX47, are targets of ERK1/2 kinases supporting this thesis^51,52^. However, other potential mechanisms also exist such as indirect modulation of ubiquitination or proteosomal activity, which can also influence SG dynamics^53–56^.

Another key finding is the pleiotropic effect of small molecules that inhibit the NF-κB signaling pathway on SG stability. Several of these compounds induced spontaneous SGs but they also destabilized SGs that were pre-formed. We found that PPM18 engages the canonical pathway of SG assembly through eIF2α phosphorylation, but whether other NF-κB inhibitors that induce SGs also act through the same pathway remains unknown. Interestingly, inhibition of IKKβ did not have an effect on SG stability suggesting that IKKα might be playing a more critical role in the crosstalk between innate immune and stress signaling pathways. It could also explain differential effects of NF-κB inhibitors because they are expected to have varying inhibition efficacy against the two kinases. Since SGs are dynamic supramolecular structures in which assembly and disassembly processes are active simultaneously, accelerating rate of disassembly can also destabilize SGs^57–59^. Alternatively, SGs are formed and get disassembled as part of their natural “*life cycle*”. NF-κB inhibitors that spontaneously induce SGs could speed up these processes ultimately resulting in faster clearance of SGs. Which of these models, or another, can explain destabilization of SGs by several NF-κB inhibitors still needs to be tested.

PPM18 induces caspase independent RCD. Several such pathways have been discovered over the last two decades^60^. A previous study had reported that PPM18 induces apoptosis in bladder cancer cells^61^. However, given that the pan-caspase inhibitor zVAD failed to stop the induction of RCD, which specific pathway is engaged by PPM18 remains an open question. A critical finding of our study is that PPM18 can spontaneously induce SGs and also inhibit viral replication. However, this inhibition was independent of SG assembly because PPM18 treatment inhibited IAV replication in *G3BP1*^-/-^ cells even though these cells did not assemble SGs. Notably, there was increased viral replication in *G3BP1*^-/-^ cells without PPM18 treatment. This indicates that G3BP1 protein does play an antiviral role during IAV infection. However, it is dispensable for PPM18 mediated viral replication inhibition. These findings raise three fundamental questions. How does G3BP1 inhibit IAV replication independent of its role as a SG scaffold protein? Are SGs even required for viral replication inhibition, or is restriction of viral replication observed in their presence due to another undiscovered phenomenon happening in the background? Several studies have reported that SGs protect cells from RCD. Then why is PPM18 equally cytotoxic for *G3BP1*^-/-^ cells? There are some hints regarding the first question. G3BP1 promotes decay of RNA molecules that contain complex secondary structures^62^. If G3BP1 preferentially promotes viral RNA decay, it can potentially explain its inhibitory effect on viral replication. The second and third questions though would require taking a step back and performing a deeper investigation of SG biology and the cellular processes that are activated, or inactivated, during its assembly and disassembly. This is especially true in light of recent studies reporting that there is a stronger induction of type I interferon signaling in the absence of PKR or G3BP1 and challenges the model where SGs were thought to be acting as a molecular platform to enhance sensing of viral PAMPs by host PRRs^27,28^.

### Conclusion

Using a pilot small-molecule screen, we identified several compounds that affect SG assembly and stability. We found that one of them, PPM18, robustly inhibited IAV replication in a SG independent manner. Our findings also raise several fundamental questions about SG biology and its role in cellular homeostasis that will need to be further investigated in the future.

## Materials and Methods

### Cell culture and stimulations

A549 cells were grown in DMEM complete (DMEM media supplemented with 10% FBS (S1620; Biowest) and 1% penicillin and streptomycin (Sigma-Aldrich)) media. Cells were seeded in optimal bottom half-area 96-well plates the day before experiments (Greiner Inc). In the morning of experiments cells were washed with warm PBS and incubated in fresh DMEM complete media for 30 minutes. Transfection solution was prepared in Xfect buffer with low-molecular weight poly(I:C) (tlrl-picw-250; InvivoGen) using manufacturer’s protocol (Xfect transfection kit, Takara Bio). In parallel, DMEM media was removed, and cells were washed again with PBS. The transfection solution was diluted with warm Opti-MEM media (ThermoFisher Scientific) so that the final concentration of poly(I:C) was 1 µg/mL. One hundred microliters of this solution was immediately added to cells and incubated for 6 hours. At 6 hours, Opti-MEM was removed and warm DMEM complete media containing inhibitors or DMSO was added to the cells. One hour after adding inhibitors, cells were fixed with 4% paraformaldehyde and processed for immunofluorescence confocal microscopy as described in the following section. Sodium arsenite at 500 µM concentration was used as previously described^31^. Propidium iodide staining based RCD measurement was performed as previously described^4^, with one modification. Images were acquired on ImageXpress Pico automated microscope. *G3BP1*^-/-^ cells were generated using a gRNA expressing previously described adenoviral plasmid that also expresses CAS9 and GFP (px458_2A_GFP_sgRNA_G3BP1_G1)^63^. For generating monoclonal cell lines, GFP positive cells were individually sorted into half-area 96-well plates and cultured for 14 days in DMEM complete media. Cells were expanded into 24-well tissue culture plates followed by 6-well tissue culture plates. Western blot analysis was used to screen for clones that lacked G3BP1 expression. Three clones with complete loss of G3BP1 protein randomly were selected for further experiments. Cells knockout for EIF2AK1, EIF2AK2, EIF2AK3, and EIF2AK4 were generated using lentiviral transduction. Two previously described gRNAs each were used^64^. After selection, transduced cells were selected three times using 5 µg/mL puromycin. Western blot analysis was used to validate gene knockouts for EIF2AK1, EIF2AK2, and EIF2AK3.

### Immunostaining and confocal microscopy imaging

Following stimulations cells were fixed in 4% paraformaldehyde (ChemCruz) at room temperature for 15 min and washed with PBS. Blocking was done in 2% BSA in PBS (Sigma-Aldrich). Cells were stained with mouse anti-G3BP1 (1:1000; 27299-I-AP; Proteintech), goat anti-Influenza A Virus polyclonal antibody (abcam, ab20841) or rabbit anti-DDX3X (1:1000; A300-474A; Bethyl Laboratories) antibody overnight at 4°C. Exact antibody multiplexing is described in figures. Following secondary antibodies were used: donkey Alexa Fluor 488–conjugated anti-mouse IgG (1:1000; A32766; Life Technologies), Donkey anti-Goat IgG (H+L) Highly Cross-Adsorbed Secondary Antibody, Alexa Fluor™ Plus 555 (1:1000; A32816; Life Technologies), and Donkey anti-Rabbit IgG (H+L) Highly Cross-Adsorbed Secondary Antibody, Alexa Fluor™ Plus 555 (1:1000; A32794; Life Technologies). DAPI (5 μg/ml, Cayman Chemicals Inc) was used to visualize nuclei. Confocal images were acquired on ImageXpress® Micro Confocal (Molecular Devices) confocal microscope. Images were analyzed as described in the following section. ImageJ Fiji was used for preparing images for publication^65^.

### Training of artificial intelligence models for image analysis and data processing

SG were detected using ilastik and CellProfiler softwares^35,36^. Initial segmentation was performed in ilastik, where a pixel classification workflow was trained to distinguish SG from the background. The probability maps generated by ilastik were then imported into CellProfiler for object identification and measurement. In CellProfiler, granules were segmented using IdentifyPrimaryObjects module based on the probability maps, and SG numbers were quantified using MeasureObjectSizeShape module. This pipeline enabled robust and automated analysis of SG dynamics across experimental conditions.

### Western blot analysis

Western blot analysis was performed as described in^9^. Primary Abs used were anti–phospho-eIF2α (1:1000; catalog no. 3398; Cell Signaling Technology [CST]), anti-eIF2α (1:1000; catalog no. 9722; CST), anti-G3BP1 (1:10000, Catalog no. 66486-1-Ig; Proteintech), anti-EIF2AK1 (1:1000, Catalog no. 20499-1-AP, Proteintech), anti-EIF2AK2 (1:1000, Catalog no. 18244-1-AP, Proteintech), anti-EIF2AK3 (1:1000, Catalog no. 20582-1-AP, Proteintech), CoralLite 555-conjugated beta actin mouse monoclonal antibody (1:3000, Catalog no. CL555-66009, Proteintech) and anti-GAPDH (1:10000, Catalog no. 60004-1-Ig; Proteintech). Secondary antibody used were Peroxidase-conjugated AffiniPure Goat Anti-Rabbit IgG, F(ab’)2 Fragment Specific (1:1000, Catalog no. 111-035-047; Jackson Immunoresearch) and Peroxidase-conjugated AffiniPure Rabbit Anti-Mouse IgG, F(ab’)2 (1:10000, Catalog no. 315-035-047; Jackson Immunoresearch).

### Reverse transcription quantitative PCR analysis

Following the stimulations, cells were lysed in TRIZOL (Thermo Fisher Scientific, 15596026) at indicated time-points followed by RNA extraction according to the manufacturer’s protocol. 250 ng of RNA was reverse transcribed using High-Capacity cDNA Reverse Transcriptase kit (Applied Biosystem, 4368813). Quantitative PCR was done on the BioRad CFX Connect^TM^ real-time PCR instrument using 2× SYBR Green (Applied Biosystems, 4368706). The primers used are as follows: *Gapdh*: 5′-CGTCCCGTAGACAAAATGGT-3′, 5′-TTGATGGCA ACAATCTCCAC-3′; IAV: 5′-ATGGATCCAAACACTGTGTC-3’, 5′-TCAAACTTCTGACCTAATTGTTCC-3’, *TNF*: 5′-CCTCTCTCTAATCAGCCCTCTG-3′, 5′-GAGGACCTGGGAGTAGATGAG-3′, *IFNB*: 5′-CAACTTGCTTGGATTCCTACAAAG-3′, 5′-TATTCAAGCCTCCCATTCAATTG-3′.

### Statistical analysis

Statistical significance of the data was determined by the two-tailed Student’s *t*-test or one-way ANOVA with alpha level of 0.05. Sample means are reported, and error bars represent SEM. GraphPad Prism v8 software was used for statistical analysis and generating plots. n.s. = not significant. \**p*-value <= 0.05, \*\**p*-value <= 0.01, \*\*\**p*-value <= 0.001, and \*\*\*\**p*-value <= 0.0001.

## Acknowledgements.

We thank all members of the Department of Microbiology and Immunology at the University of Texas Medical Branch for their advice and encouragement. We would especially like to thank Dr. Ashok Chopra, Dr. Janice Endsley, Dr. Alexander Freiberg, Dr. Matthieu Gagnon, Dr. Lynn Soong, and Dr. Scott Weaver for their mentorship. px458_2A_GFP_sgRNA_G3BP1_G1 was a gift from Thomas Tuschl (Addgene plasmid # 127114; http://n2t.net/addgene:127114 ; RRID:Addgene_127114). EIF2AK1 gRNA (BRDN0001147917), EIF2AK2 gRNA (BRDN0001146342), EIF2AK3 gRNA (BRDN0001145848), and EIF2AK4 gRNA (BRDN0001145948) were gifts from John Doench & David Root (Addgene plasmid #’s 77047, 75637, 77166, and 75875).

## Author contributions

PS designed the experiments. PPL, A, and PS performed the experiments. XX and PS supervised the study. PS and A wrote the first draft of the manuscript. All authors contributed to writing of the manuscript.

---

**Supplementary figure 1:**
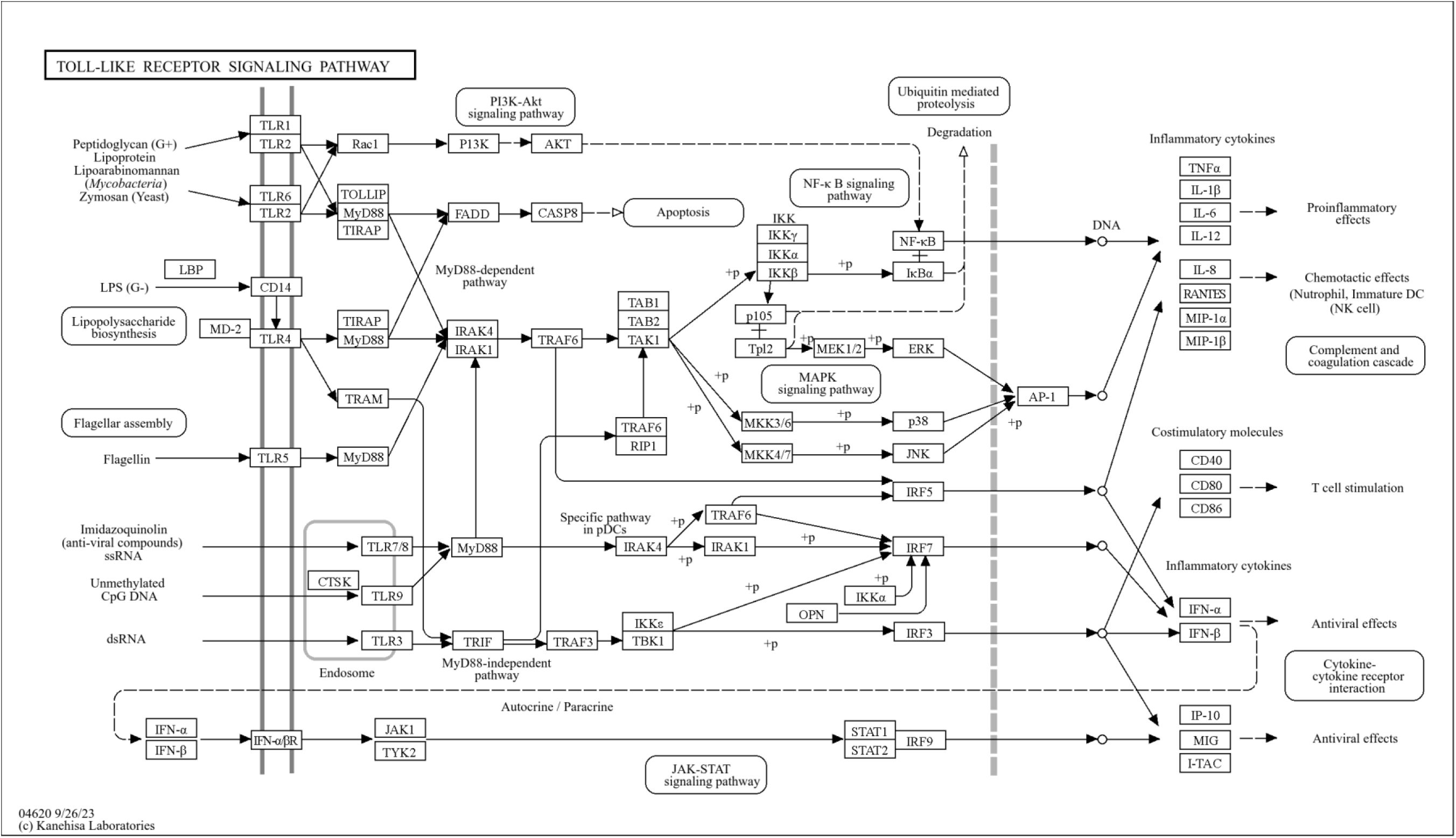
Network map of the toll-like signaling network in the Kyoto Encyclopedia of Genes and Genomes (KEGG) database. The network map was downloaded from the KEGG database for visualizing the connections in the TLR signaling pathway. Empirically selected kinases in the TLR signaling pathway were targeted for inhibition in the pilot screen reported in this manuscript.

**Supplementary table 1: List of inhibitors, their targets and concentrations.**

